# Environmental DNA Metabarcoding for Simultaneous Monitoring and Ecological Assessment of Many Harmful Algae

**DOI:** 10.1101/2020.10.01.322941

**Authors:** Emily Jacobs-Palmer, Ramón Gallego, Kelly Cribari, Abigail Keller, Ryan P. Kelly

**Affiliations:** University of Washington School of Marine and Environmental Affairs; NRC Research Associateship Program, Northwest Fisheries Science Center, National Marine Fisheries Service, National Oceanic and Atmospheric Administration.

## Abstract

Harmful algae can have profound economic, environmental, and social consequences. As the timing, frequency, and severity of harmful algal blooms (HABs) change alongside global climate, efficient tools to monitor and understand the current ecological context of these taxa are increasingly important. Here we employ environmental DNA metabarcoding to identify patterns in a wide variety of harmful algae and associated ecological communities in the Hood Canal of Puget Sound in Washington State, USA. We track trends of presence and abundance in a series of water samples across nearly two years. We find putative harmful algal sequences in a majority of samples, suggesting that these groups are routinely present in local waters. We report patterns in variants of the economically important genus *Pseudo-nitzschia* (family Bacillariaceae), as well as multiple harmful algal taxa previously unknown or poorly documented in the region, including a cold-water variant from the saxitoxin-producing genus *Alexandrium* (family Gonyaulacaceae), two variants from the karlotoxin-producing genus *Karlodinium* (family Kareniaceae), and one variant from the parasitic genus *Hematodinium* (family Syndiniaceae). We then use data on environmental variables and the biological community surrounding each algal taxon to illustrate the ecological context in which these species are commonly found. Environmental DNA metabarcoding thus simultaneously (1) alerts us to potential new or cryptic occurrences of harmful algae, (2) expands our knowledge of the co-occurring conditions and species associated with the growth of these organisms in changing marine environments, and (3) provides a tool for monitoring and management moving forward.

## Introduction

Harmful algae and associated blooms create environmental, health, and economic challenges at a global scale, causing mass die-offs in ecosystems from de-oxygenation (Gobler, 2020; Griffith & Gobler, 2020), multiple types of poisoning in humans (Trainer et al., 2013), and significant losses of revenue for the aquaculture industry (Trainer & Yoshida, 2014; Dìaz et al., 2019). For these reasons, local, national, and international governing bodies organize and fund monitoring programs to track HABs and identify the conditions that lead to their occurrence (Graneli & Lipiatou, 2002; Trainer, 2002; Lopez et al., 2008; Moestrup et al., 2020). In addition, changing marine environments appear to be causing increases in the duration, frequency, and severity of HABs globally in association with rising temperatures and declining pH (Gattuso et al., 2015a; Gobler et al., 2017).

Hood Canal, a natural glacial fjord within the Puget Sound of Washington, USA, is a useful natural system in which to study the ecology of harmful algae and likely future changes to their patterns of occurrence. Surface temperatures of the region have risen 1.0°C since the 1950s, dissolved oxygen levels are below 5 mg/L in deeper sections of the sound, and pH has dropped by 0.05 - 0.15 units since pre-industrial era (~1750) (Feely et al., 2010; Busch, Harvey & McElhany, 2013; Mauger et al., 2015). Warmer temperatures and longer durations of warm conditions will create larger windows of growth for some HABs moving forward (Moore, Mantua & Salathe Jr, 2011; Mauger et al., 2015), with ocean acidification exacerbating the impacts of these blooms by further increasing the toxicity and growth of harmful algal species (Fu, Tatters & Hutchins, 2012; Field et al., 2014).

Harmful algae fall into four primary categories: diatoms, dinoflagellates, haptophytes, and raphidophytes. Of particular concern locally are diatoms from the genus *Pseudo-nitzschia* and dinoflagellates from the genera *Alexandrium*, *Gonyaulax*, and *Protoceratium*, each of which produces toxins that can accumulate in shellfish grazers (Shimizu et al., 1975; Satake, MacKenzie & Yasumoto, 1998; Cembella, Lewis & Quilliam, 2000; Trainer et al., 2009, 2016). When consumed by humans, the toxins then cause symptoms ranging from amnesia to paralysis, and can be deadly (Ferrante et al., 2013; Grattan, Holobaugh & Morris Jr, 2016). Additional harmful alga of concern are fish-killing species such as the diatom *Chaetoceros concavicornis* and the raphidophyte *Heterosigma akashiwo* (Yang & Albright, 1994; Khan, Arakawa & Onoue, 1997). There is no current ensemble testing protocol for all of these local problematic algae, and both human-mediated transport and warming-related range shifts are likely to introduce additional taxa. For example, there is recent evidence that the toxic dinoflagellate *Karenia mikimotoi* (family Kareniaceae; described from Japan and also occurring in the North Atlantic) is now present along the west coast of North America, specifically off of Alaska and California (National Centers for Coastal Ocean Science, 2014).

Because the effects of harmful algae are wide-ranging and potentially devastating (Lewitus et al., 2012; Moore et al., 2019), monitoring these organisms and the environmental conditions with which they are associated has long been a public health priority in Puget Sound and in many locations around the world. Efforts to track blooms of harmful algae have historically relied on the work of skilled taxonomists using microscopic visual analysis of cells to identify species (e.g. Lapworth, Hallegraeff & Ajani, 2001; Yang et al., 2000). More recently, satellite spectrographic data (Tomlinson et al., 2004; Ahn et al., 2006), molecular assays for toxins (Pierce & Kirkpatrick, 2001; Murray et al., 2011), and flow cytometry coupled with machine learning (Campbell et al., 2010) have been employed to detect and track HABs.

Adding to the list of technological advances for monitoring are two types of genetic techniques that rely on environmental DNA (eDNA) present in the water to classify and assess the abundance of harmful algae: quantitative Polymerase Chain Reaction (qPCR) and DNA metabarcoding (Al-Tebrineh et al., 2010; Erdner et al., 2010; Antonella & Luca, 2013; Grzebyk et al., 2017; Ruvindy et al., 2018). The former method tracks known taxa individually and requires substantial sequence data to design species-specific primers and/or probes; the latter method involves PCR with less-specific primers to generate amplicons from a broad swath of taxa at a common locus. This second method, metabarcoding, allows detection and even quantification of many taxa simultaneously.

Because the specific target organisms need not be chosen *a priori*, metabarcoding may uncover taxa unexpected in the study region, and in addition, can reveal a cross-section of the biological community surrounding any particular group of interest (Deiner et al., 2017; Taberlet et al., 2018). Previous work has described the ecological context of harmful algae and their blooms using environmental covariates (e.g. Wells et al., 2015; Banerji et al., 2019), as well as assessments of bloom-associated taxa (typically other microorganisms or viruses) (e.g. Loureiro et al., 2011; Buskey, 2008), although the labor required for traditional survey methodology has limited the breadth of contextual taxa included in these studies.

Here, we couple environmental monitoring with eDNA metabarcoding to track the presence of dozens of potential harmful algal taxa simultaneously, including a number unexpected or understudied in the region of interest. Because of their economic and public-health importance, we focus on variants in the genera *Alexandrium* (dinoflagellate family Gonyaulacaceae), *Hematodinium* (dinoflagellate family Syndiniaceae), *Karlodinium* (dinoflagellate family Kareniaceae), and *Pseudo-nitzschia* (diatom family Bacillariaceae), and examine the distribution of these in time and space across 19 months of sampling at ten locations in Hood Canal. We model the associations of individual algal lineages with key environmental variables, and subsequently improve the predictive value of these models by including co-occurring (non-algal) taxa as possible indicator species. With these analyses, we present useful information on the distributions and ecological contexts of potentially harmful algae in the Puget Sound region, and demonstrate how eDNA metabarcoding can improve our understanding and management of harmful algae, both locally and globally.

## Materials and Methods

### Environmental DNA Sampling and Measuring Environmental Variables

To identify a broad range of potential harmful algal taxa and simultaneously survey the surrounding biological community, we sampled seawater for eDNA from ten sites within Hood Canal, a natural glacial fjord in Puget Sound, Washington, USA. Five sampling sites were intertidal, and five were nearby nearshore locations at the approximate center of the fjord. For intertidal locations at Salisbury Point County Park (SA), Triton Cove State Park (TR), Lilliwaup Tidelands State Park (LL), Potlatch State Park (PO), and Twanoh State Park (TW) (see Figure 1 and Table S1 for location coordinates), we collected three 1 L samples of water from immediately below the surface using a bleach-cleaned plastic bottle held at the end of a 1.7 meter pole. We sampled intertidal locations every 1-2 months between March 2017 and July 2018 (see Table S1 for sampling dates). At these same stations and simultaneous with eDNA sampling, we collected one 120 ml water sample from each site and poisoned it with 0.1 ml of saturated HgCl_2_ for carbonate chemistry analysis including pH (Dickson, Sabine & Christian, 2007). We also collected in situ measurements of temperature and salinity using a handheld multiprobe (Hanna Instruments, USA) and a portable refractometer. We characterized sample carbonate chemistry by measuring Total Alkalinity (TA; open-cell automated titration based on a Dosimat plus (Metrohm AG) as part of a custom system assembled by Andrew Dickson (University of California San Diego) and used in the laboratory of Alex Gagnon at the University of Washington) and Dissolved Inorganic Carbon (DIC; Apollo Instruments, USA; CO2 extraction system with 10% (v/v) phosphoric acid). Both measurements were calibrated and validated with certified reference material from the Scripps Oceanographic Institute. Using DIC and TA, we calculated pH and the remaining carbonate system parameters with the R package ‘seacarb’ (Gattuso et al., 2015b), removing a single outlier sample from the dataset used for environmental modeling (see below) due to an unreasonably low pH value (<7.5).

**Figure 1:**
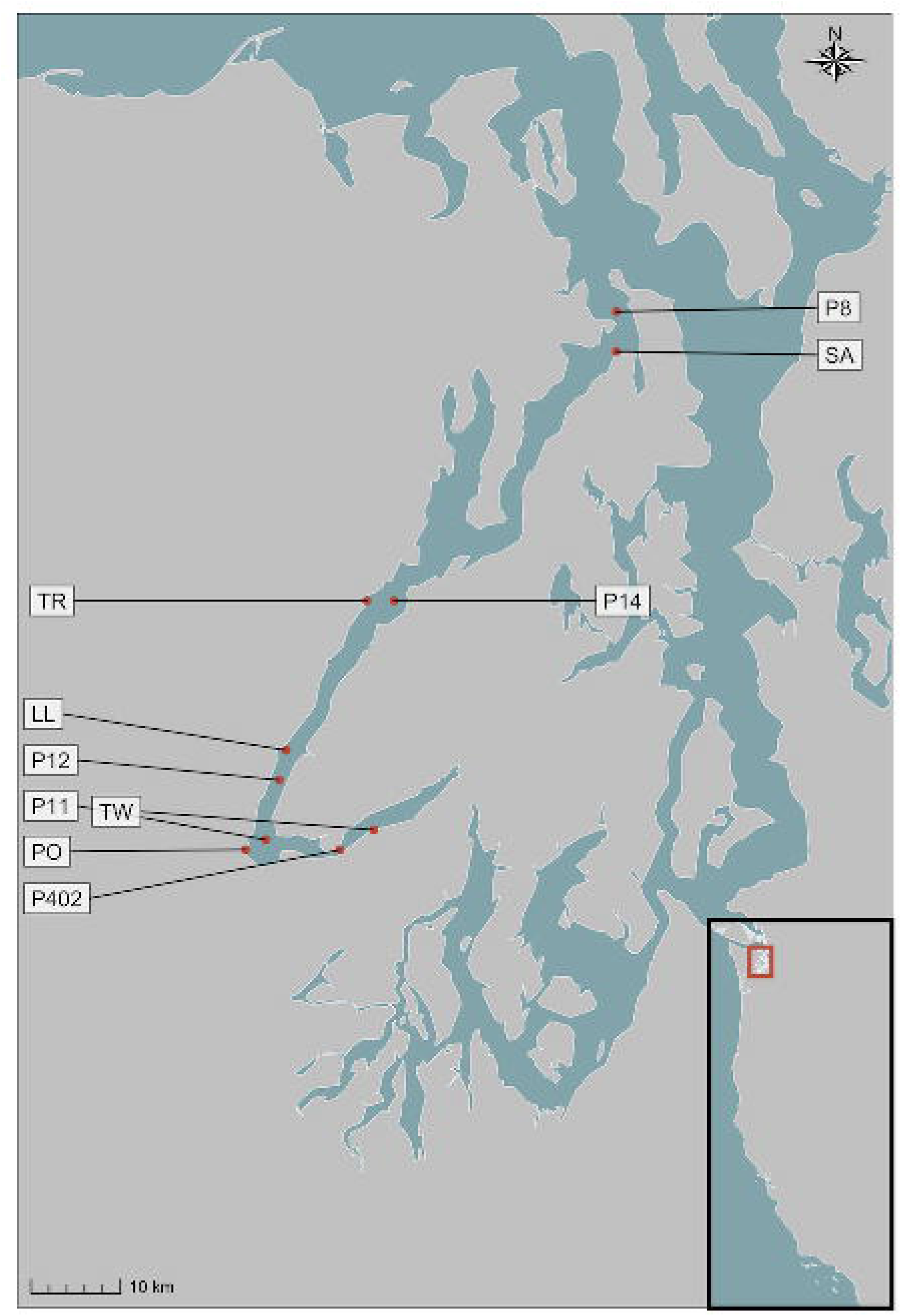
Intertidal and nearshore sampling locations in Hood Canal, Washington, USA. Site abbreviations are described in the text, and coordinates are given in Table S1. Inset map shows the Pacific coast of the continental United States.

For nearshore locations, we sampled a selection of stations (P8, P14, P12, P11, P402) surveyed by the Washington Ocean Acidification Center during triannual cruises (see Figure 1 and Table S1 for location coordinates). The samples used here were collected in September 2017 (2 samples), April 2018 (3 samples), and September 2018 (3 samples); see Table S1 for sampling dates. At each station, a CTD was deployed with twelve Niskin bottles, and collected data on temperature, salinity, and pH (Alin et al., 2019a,b,c) in addition to water for eDNA from immediately below the surface. We filtered 500 mL of each water sample for eDNA from both intertidal and nearshore locations with a cellulose acetate filter (47 mm diameter, 0.45 µm pore size), and preserved this filter in Longmire buffer until DNA extraction (Renshaw et al., 2015). Many unmeasured variables influence planktonic communities (e.g., nutrients, sunlight, and wave energy); nevertheless the minimal set of parameters we analyzed here clearly distinguished communities and was adequate for the purposes of assessing temporal and spatial trends. Our purpose was to describe patterns of harmful algae over space and time, along with the environmental and ecological contexts in which they occurred, rather than to test any particular mechanism by which harmful algal taxa might respond to different environmental parameters.

### Extraction, Amplification, and Sequencing

To extract DNA from sample filters, we used a phenol:chloroform:isoamyl alcohol protocol (modified from Renshaw et al., 2015). To maximize extraction efficiency and minimize co-extraction of inhibitors, we incubated filter membranes at 56° C for 30 min before adding 900 µL of phenol:chloroform:isoamyl alcohol and shaking vigorously for 60 s. We conducted two consecutive chloroform washes by centrifuging at 14,000 rpm for 5 min, transferring the aqueous layer to 700 µL chloroform, and shaking vigorously for 60 s. After a third centrifugation, we transferred 500 µL of the aqueous layer to tubes containing 20 µL 5 molar NaCl and 500 µL 100% isopropanol, and froze these at −20° C for approximately 15 h. Finally, we centrifuged samples at 14,000 rpm for 10 min, poured off or pipetted out any remaining liquid, and dried in a vacuum centrifuge at 45° C for 15 min. We resuspended the eluate in 200 µL water, and used 1 µL of diluted DNA extract (between 1:10 and 1:400) as template for PCR.

To identify a wide variety of metazoan taxa including putative harmful algae and their surrounding biological communities from eDNA, we amplified a ~315 base pair segment of the Cytochrome Oxidase I (COI) using universal primers described in Leray et al. (2013). To distinguish technical from biological variance and to quantify each, we ran and sequenced in triplicate PCR reactions from each of the samples (i.e., individual bottles of water). For multiplex sequencing on an Illumina MiSeq, we followed a two-step PCR protocol (O’Donnell et al., 2016) with redundant 3’ and 5’ indexing. In the first step, we used a PCR reaction containing 1X HotStar Buffer, 2.5 mM MgCl2, 0.5 mM dNTP, 0.3 µM of each primer, and 0.5 units of HotStar Taq (Qiagen Corp., Valencia, CA, USA) per 20 µL reaction. The PCR protocol for this step consisted of 40 cycles, including an annealing touchdown from 62° C to 46° C (−1° C per cycle), followed by 25 cycles at 46° C. In the second step, we added identical 6 base-pair nucleotide tags to both ends of our amplicons, with unique index sequences for each individual PCR reaction. We allowed for no sequencing error in these tags; only sequences with identical tags on both the forward and reverse read-directions survived quality control. This gave us high confidence in assigning amplicons back to individual field samples.

We generated amplicons with the same replication scheme for positive control kangaroo (genus *Macropus*) tissue, selected because this genus is absent from the sampling sites and common molecular biology reagents, but amplifies well with the universal primer set used in this study. We could therefore use positive control samples to identify possible cross-contamination: reads from other taxa that appear in these samples allow us to estimate and account for the proportion of sequences that are present in the incorrect PCR reaction (see Bioinformatics below). We also amplified negative controls (molecular grade water) in triplicate alongside environmental samples and positive controls, and verified by gel electrophoresis and fluorometry that these PCR reactions contained no appreciable amount of DNA (see Kelly, Gallego & Jacobs-Palmer, 2018 for a discussion of the merits of sequencing positive and not negative controls).

To prepare libraries of replicated, indexed samples and positive controls, we followed manufacturers’ protocols (KAPA Biosystems, Wilmington, MA, USA; NEXTflex DNA barcodes; BIOO Scientific, Austin, TX, USA). We then performed sequencing on an Illumina MiSeq (250-300 bp, paired-end) platform in seven different sets of samples: six for the intertidal dataset and one for the nearshore dataset.

### Bioinformatics

We followed updated versions of previously published procedures for bioinformatics, quality control, and decontamination (Kelly, Gallego & Jacobs-Palmer, 2018). This protocol uses a custom Unix-based script (Gallego) calling third-party programs to perform initial quality control on sequence reads from all four runs combined, demultiplexing sequences to their sample of origin and clustering unique variants into amplicon sequence variants (ASVs) (Martin, 2011; Callahan et al., 2016).

Specifically, to address possible cross-sample contamination (see Schnell, Bohmann & Gilbert, 2015), we subtracted the maximum proportional representation of each ASV across all positive control samples (see Extraction, Amplification, and Sequencing above) from the respective ASV in field samples. We estimated the probability of ASV occurrence by performing occupancy modeling (Royle & Link, 2006; Lahoz-Monfort, Guillera-Arroita & Tingley, 2016). Following Lahoz-Monfort et al. (2016) and using the full Bayesian code for package rjags (Plummer et al., 2016) provided by those authors, we modeled the probability of occupancy (i.e., true presence) for each of the unique sequence variants in our dataset. We treated replicate PCR reactions of each water bottle as independent trials, estimating the true-positive rate of detection (P11), false-positive rate (P_10_), and commonness (psi, 1/) in a binomial model. We then used these parameters to estimate the overall likelihood of occupancy (true presence) for each ASV; those with low likelihoods (<80%) were deemed unlikely to be truly present in the dataset, and therefore culled. We removed samples whose PCR replicates were highly dissimilar by calculating the Bray-Curtis dissimilarity amongst PCR replicates from the same bottle of water and discarding those with distance to the sample centroid outside a 95% confidence interval. The result was a dataset of 3.98 × 10^8^ reads from 5275 unique ASVs. Lastly, to collapse variants likely due to PCR error, we converted ASVs to operational taxonomic units (OTUs) by clustering with SWARM (Mahé et al., 2015). All bioinformatic and analytical code is included in a GitHub repository (https://github.com/ramongallego/Harmful.Algae.eDNA), including the details of parameter settings in the bioinformatics pipelines used. Sequence and annotation information are included as well, and the former are deposited and publicly available in GenBank (upon acceptance; accession numbers will be provided in the published manuscript).

### Taxonomy

We performed the taxonomic identification using a CRUX-generated database for the Leray fragment of the COI gene (see Extraction, Amplification, and Sequencing above), querying that database with a Bowtie2 algorithm (as described in Curd et al., 2019). The algorithm classifies the query sequence to the last common ancestor of ambiguously classified sequences. Only matches with a bootstrap support greater than 90% were kept. Here, we assigned taxonomy at the level of genus, rather than species, for two main reasons. First, for some taxa, variation may not be sufficient to distinguish species within a genus, and second, representation of local species in the databases used may not be complete, leading to the mis-assignment of sequences to their nearest represented neighbor. We denoted different lineages within genera using three-character abbreviation derived from the sequence variants themselves. Full sequences for each variant are provided in Table S2. To assess similarity of putative harmful algal lineages, we translated nucleotide sequences with the ExPASy Bioinformatics Resource Portal Translate tool using the mold, protozoan, and coelenterate mitochondrial, mycoplasma/spiroplasma genetic code (Gasteiger et al., 2003). We created both nucleotide and amino acid alignments with the Clustal Omega Multiple Sequence Alignment tool (Sievers & Higgins, 2014).

### Species Distributions in Space and Time

To examine the distribution of potential harmful algal taxa in time and space, we calculated an index of relative eDNA abundance (hereafter eDNA abundance index). To derive this index, we first normalized taxon-specific ASV counts into proportions within a technical replicate, and then transformed the proportion values such that the maximum across all samples was scaled to 1 for each taxon (Kelly, Shelton & Gallego, 2019). Such indexing allowed us to track trends in abundance of taxa in time and space by correcting for both differences in read depth among samples and differences in amplification efficiency among sequences. We plotted the eDNA abundance index for each potential harmful algal taxon across all sampling events from both intertidal and nearshore eDNA collections in our time series between March 2017 and September 2018.

### Environmental and Biological Context

To explore the ways in which environmental variables were associated with the presence or absence of our focal harmful algal taxa, we compared logistic-regression models using taxon presence as outcome, and combinations of three environmental variables (temperature, pH, and salinity) as predictors. We also fit a variety of models in a Bayesian hierarchical framework, where the slopes of predictors and intercepts could vary by season (summer/winter), and included all models (hierarchical and non-hierarchical) in our model comparison. Rather than mechanically testing all possible combinations of models, we proposed models that were reasonable given the observed patterns of occurrences; in total, this resulted in between five and 17 models per taxon. For purposes of the models, we designated April through September as being “summer”, and other months “winter”. Given many possible predictor variables, developing a useful model without overfitting can be a challenge. To combat this, we compared models using the widely applicable information criterion (WAIC) (Watanabe, 2010), which makes no assumptions about the shape of the posterior probability distribution and – like information criteria in general – penalizes more complex models. Moreover, WAIC quickly approximates the results of leave-one-out cross-validation (McElreath, 2020) to estimate out-of-sample model performance. Following model selection using WAIC, we reported the in-sample model accuracy for reference.

To determine the species most closely associated with potential harmful algal taxa, we performed canonical analysis of principal coordinates (CAP) for each focal variant by implementing the capscale function in vegan (Oksanen et al., 2013), which revealed the degree to which other taxa in the surrounding biological communities could be associated with presence (versus absence) of the potential harmful algal taxon. Using this ordination technique avoids the problem of testing each co-occurring taxon for significant associations with our focal putative harmful algae, thereby removing the need to statistically correct for multiple comparisons. We then used these putative indicator taxa as predictors in a second round of logistic regressions, adding only the single most-strongly associated taxon as a separate intercept term to the best-fit environmental models for each of our focal lineages (above). Such contextual ecological information is useful to the extent that it helps to predict the occurrence of harmful algal lineages without overfitting, which we evaluated as described above using WAIC.

## Results

### Taxonomy

Environmental DNA metabarcoding of 63 samples from five intertidal and five nearshore locations in Hood Canal, Washington, United States revealed a total of 605 unique amplicon sequence variants (ASVs) for which we were able to assign taxonomy to 262 distinct genera. Of these, exactly 100 ASVs were assigned to genera that are known to contain harmful algae (Horner & Postel, 1993; Horner, Garrison & Plumley, 1997; Trainer et al., 2016; Moestrup et al., 2020). These potential harmful algal taxa are members of four main taxonomic groups – diatoms, dinoflagellates, haptophytes, and raphidophytes – and represent seventeen genera (diatoms: *Chaetoceros*, *Nitzschia*, *Pseudo-nitzschia*; dinoflagellates: *Alexandrium*, *Dinophysis*, *Gonyaulax*, *Gymnodinium (Akashiwo)*, *Hematodinium*, *Heterocapsa*, *Karlodinium*, *Prorocentrum*, *Woloszynskia*; haptophytes: *Chrysocromulina*, *Phaeocystis*; raphidophytes: *Chattonella*, *Heterosigma*, *Pseudochattonella*; See Table S2 for a complete list of potential harmful algal taxonomic assignments and COI sequences).

Taxa that occur only in small numbers of samples lack sufficient observations to allow robust tests for association with environmental variables. Consequently, we focus hereafter on the ASVs present in at least ten percent of samples (minimum 7 occurrences out of 63 samples), an adequate sample size to compare with environmental variables and biological context. This subset of sequences included 191 total variants, 37 of which belong to potentially harmful algal taxa (Table 1), and the rest to other members of the biological community. These putative harmful algal variants belong to 12 genera containing differing degrees of sequence variation, with some such as *Hematodinium* represented by a single DNA and protein sequence, and others such as *Nitzschia* represented by a much larger number of DNA (10) and amino acid (5) variants. Each of the potential harmful algal genera represented here exhibit varying degrees and types of toxicity or harm, ranging from physical irritation of fish gill tissue to production of toxins dangerous to human health (Table 1; Trainer et al., 2016; Simonsen & Moestrup, 1997; Lindberg, Moestrup & Daugbjerg, 2005; Stentiford & Shields, 2005; Kotaki et al., 2006; Peperzak & Poelman, 2008; Skjelbred et al., 2011; Place et al., 2012; Cho et al., 2017).

**Table 1:** Potential harmful algal taxa identified by eDNA in at least ten percent of samples from in Hood Canal, WA. Type of harmful algae and genus are given, as well as the number of DNA and protein variants, toxicity, and sampling location(s) for member(s) of that genus.

| Type | Genus | DNA variants | Protein variants | Toxicity (target) | Sampling Location |
| --- | --- | --- | --- | --- | --- |
| Diatom | Chaetoceros | 8 | 5 | Gill irritation (finfish) | Intertidal; Nearshore |
| Diatom | Nitzschia | 10 | 5 | Domoic acid and derivatives (human) | Intertidal |
| Diatom | Pseudo-nitzschia | 3 | 2 | Domoic Acid (human) | Intertidal; Nearshore |
| Dinoflagellate | Alexandrium | 2 | 2 | Saxitoxin (human) | Intertidal; Nearshore |
| Dinoflagellate | Hematodinium | 1 | 1 | Parasitism (crab) | Intertidal |
| Dinoflagellate | Heterocapsa | 2 | 2 | Haemolysis (shellfish) | Intertidal; Nearshore |
| Dinoflagellate | Karlodinium | 2 | 2 | Karlotoxin (human) | Intertidal; Nearshore |
| Dinoflagellate | Woloszynskia | 1 | 1 | Reddening of water (general) | Intertidal |
| Haptophyte | Chrysochromulina | 3 | 3 | Haemolysis (shellfish) | Intertidal; Nearshore |
| Haptophyte | Phaeocystis | 2 | 2 | Oxygen depletion (general) | Intertidal; Nearshore |
| Raphidophyte | Chattonella | 2 | 1 | Reactive oxygen species (finfish) | Intertidal; Nearshore |
| Raphidophyte | Pseudochattonella | 1 | 1 | Gill irritation (finfish) | Intertidal |

Amplicon sequences from environmental samples cannot be matched directly with phenotypes, by definition, and taxonomic annotations of those sequences depend upon adequate reference material. Acknowledging both the intra-specific variation that exists at the COI locus and the incompleteness of the GenBank reference database for many of these groups, we treat polymorphism within a putative genus as being ambiguous: these variants may be intra-specific, or they may represent distinct evolutionary lineages. For these reasons, we conservatively perform analyses on the sequence variants themselves (denoted with their genus names and a three-character code that abbreviates the hash of the unique nucleotide sequence) rather than making assumptions regarding their status as haplotypes versus species.

Putative harmful algal taxa from a few genera are of particular interest, due to the nature of their toxicity (*Alexandrium*), to their unexpected presence in the study region (*Hematodinium* and *Karlodinium*), or to their potential economic impact (*Pseudo-nitzschia*). For these reasons, we chose to examine aspects of their taxonomy, distribution, and ecology in greater detail. We first examined COI sequences for these taxa from our original metabarcoding effort, noting that both *Alexandrium* and *Karlodinium* genera were each represented by two sequence variants, *Pseudo-nitzschia* by three sequence variants, and *Hematodinium* by a single sequence variant (Table 1). Amino acid translation revealed that the two *Alexandrium* ASVs differed by a single amino acid substitution, the two *Karlodinium* ASVs differed by five substitutions, and although two of the three *Pseudo-nitzschia* sequences (Pseudonitzschia_4e5 and Pseudonitzschia_d36) were identical in amino acid sequence, they differed from the third (Pseudonitzschia_d40) by two substitutions. The results below focus on these eight sequence variants, which we hereafter refer to as our “focal lineages.”

### Species Distributions in Space and Time

To identify the seasonal and spatial distributions of taxa from our eight focal lineages, we next visualized their patterns of presence and absence in time and space (Figure 2). The variants assigned to *Alexandrium*, Alexandrium_3fc and Alexandrium_2b2, had completely non-overlapping distributions in space and time, never appearing in the same sampling event. Alexandrium_3fc appeared solely in the summer (April-September) months (25 of 43 summer samples vs. 0 of 20 winter samples; p < 0.001) whereas Alexandrium_2b2 appeared primarily in the winter (October-March) months (1 of 43 summer samples vs. 7 of 20 winter samples; p < 0.001). In contrast, the single variant assigned to *Hematodinium*, Hematodinium_449, was not significantly seasonal (9 of 43 summer samples and 3 of 20 in winter; p = 0.742); neither were the two variants assigned to *Karlodinium*, Karlodinium_8ed and Karlodinium_a27 (Karlodinium_8ed: 14 of 43 summer samples and 7 of 20 winter samples, p = 1; Karlodinium_a27: 6 of 43 summer samples and 5 of 20 winter samples, p = 0.322). One of the three variants assigned to *Pseudo-nitzschia* occurred significantly more frequently in summer than in winter months (Pseudonitzchia_d36: 10 of 43 summer samples and 0 of 20 winter samples, p = 0.023), while the others did not (Pseudonitzchia_4e5: 8 of 43 summer samples and 1 of 20 winter samples, p = 0.244; Pseudonitzchia_d40, 7 of 43 summer samples and 4 of 20 in winter; p = 0.737).

**Figure 2:**
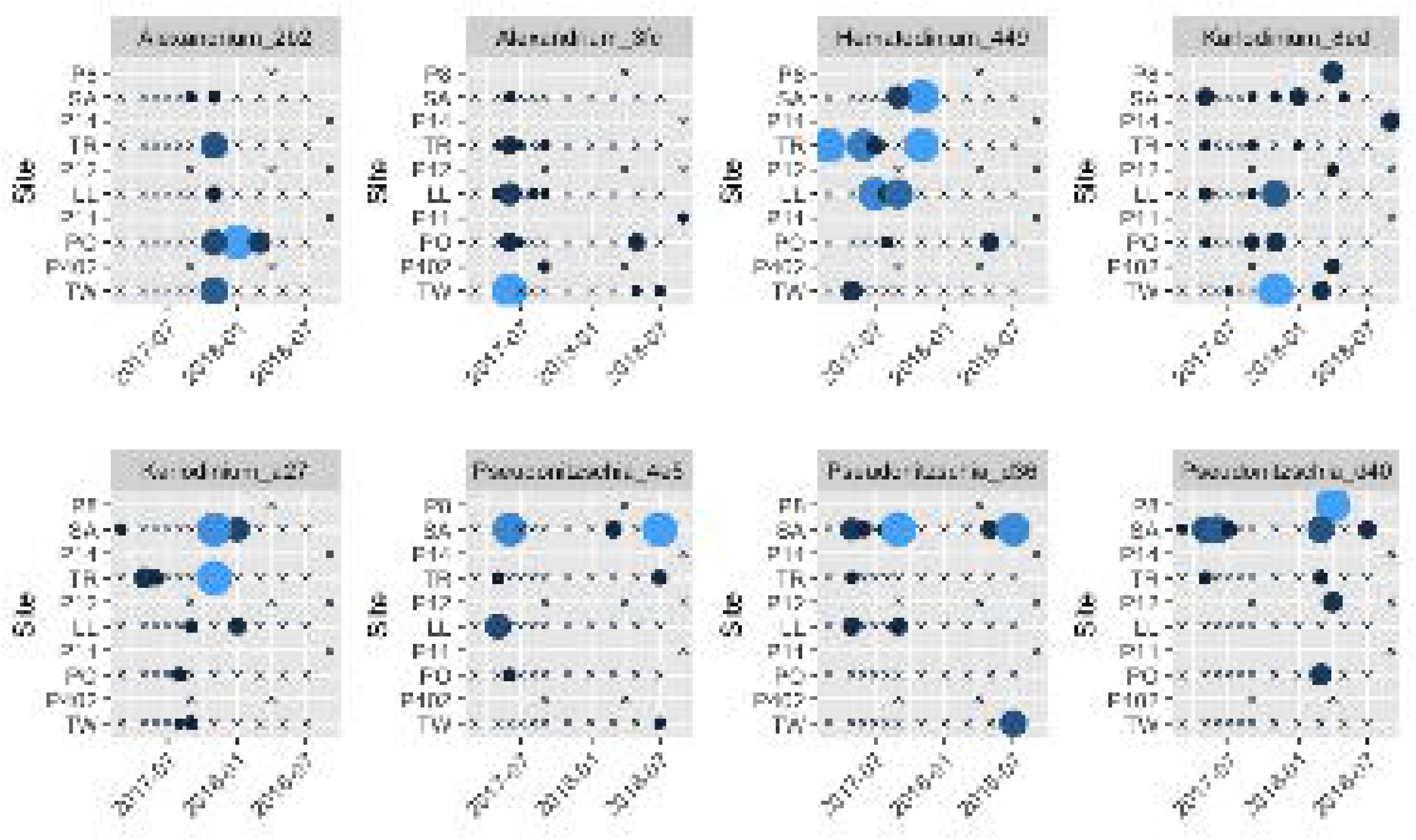
Spatial and temporal distribution of HAB taxa. Distribution of eight focal algal lineages across time and space. Larger and lighter circles indicate greater relative abundances; ‘x’ symbol indicates non-detection at that site/date. Site abbreviations are as in Figure 1 and Table S1.

All together, we detected at least one of the eight focal sequence variants in 51 out of 63 sampling events (81%), indicating that these potential harmful algal taxa are present at some level more often than not in local waters. Additionally, the larger intertidal dataset was more diverse, containing all eight focal lineages, while only three were detected on the nearshore cruises (Alexandrium_3fc, Karlodinium_8ed, Pseudonitzschia_d40).

### Environmental Context

The above results suggest both *Alexandrium* lineages and at least one of the three *Pseudo-nitzschia* lineages are associated with environmental conditions that change seasonally, while the others are more stochastic in space and time. For each focal taxon, we fit a series of logistic-regression models (see Methods) describing taxon occurrence as a function of sea-surface temperature, pH, and salinity, both with and without a global intercept term (see Table S3 for a complete list of models tested, by taxon). A subset of our models was also hierarchical, allowing slopes to vary according to season, and we used WAIC to identify the best-fit models of those tested (Table 2).

**Table 2:** Best-fit models of environmental covariates for eight focal algal lineages.

| Taxon | Environmental Model | Accuracy | True Positive Rate | True Negative Rate |
| --- | --- | --- | --- | --- |
| Alexandrium_2b2 | Intercept + pH + (1 + Temperature \| Season) | 0.92 | 0.29 | 1 |
| Alexandrium_3fc | Intercept + (1 + Temperature \| Season) | 0.69 | 0.33 | 0.84 |
| Hematodinium_449 | pH + Temperature | 0.79 | 0 | 0.98 |
| Karlodinium_8ed | Intercept + (Intercept + Temperature \| Season) | 0.76 | 0.45 | 0.9 |
| Karlodinium_a27 | pH + (Intercept + Salinity \| Season) | 0.84 | 0.09 | 1 |
| Pseudonitzschia_4e5 | Salinity + Temperature + pH | 0.87 | 0.22 | 0.98 |
| Pseudonitzschia_d36 | Intercept + Salinity + pH + (1 \| Season) | 0.89 | 0.4 | 0.98 |
| Pseudonitzschia_d40 | Salinity | 0.82 | 0.27 | 0.94 |

All but one of these models involves multiple environmental parameters, making them difficult to adequately visualize in two dimensions. Nevertheless, plotting the probability of taxon presence as a function of the single most-influential environmental variable and capturing seasonal variation in slope when models are hierarchical illustrates the degree to which the models do (or do not) explain the observed variance in potential harmful algal taxa (Figure 3). Among the environmental variables measured, both putative harmful algal variants most closely-associated with pH occur more frequently in our samples at lower, more acidic values (Hematodinium_449, and Karlodinium_a27). For those that are most closely-associated with temperature, warmer waters see higher frequencies of a majority of putative harmful algae during the season in which they primarily occur (Alexandrium_2b2 in winter, Alexandrium_3fc in summer) with the notable exception of Karlodinium_8ed, which occurs frequently year-round and shows a conflicting relationship with temperature across seasons.

Although environmental covariates sea-surface temperature, pH, and salinity are associated with the presence or absence of our eight focal lineages, accuracy of these models varies widely, from 0.69 for Alexandrium_3fc to 0.92 for Alexandrium_2b2 (Table 3). Additionally, these covariates alone can predict only a minority of occurrences for all taxa (Table 3, true positive rate). Consequently, we use eDNA metabarcoding data from the communities surrounding our focal lineages for potentially helpful information about the ecology of these putative harmful algae.

**Table 3:**
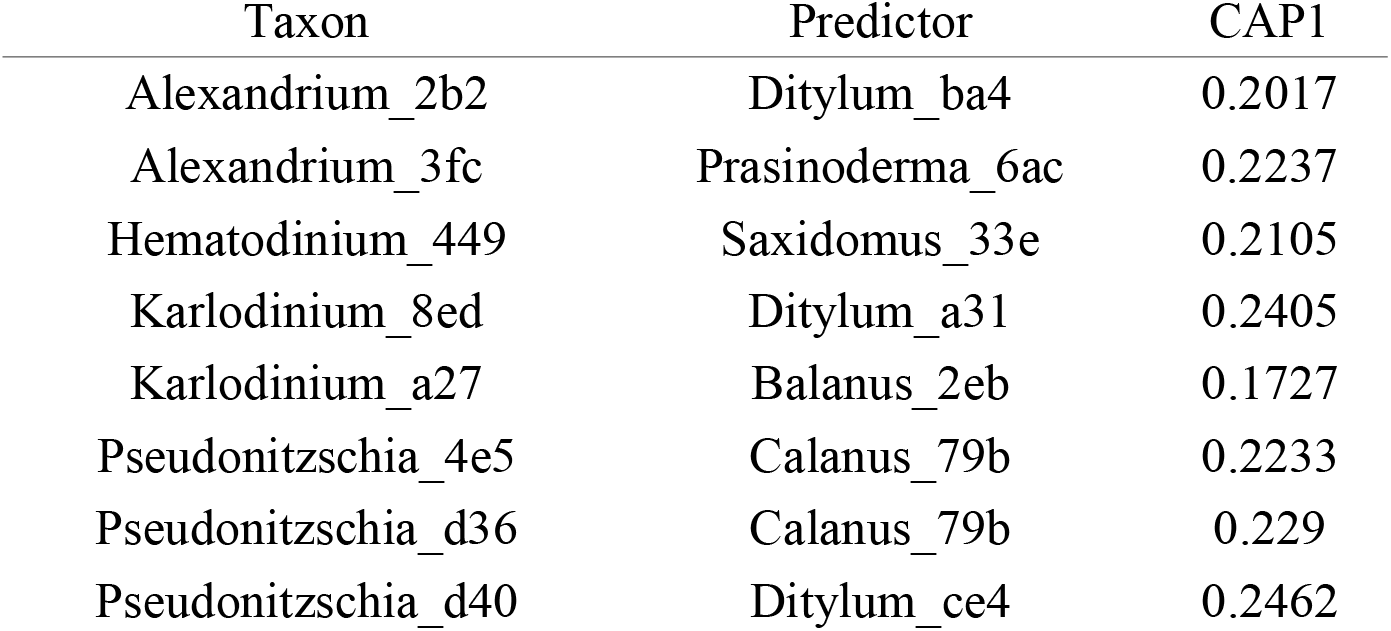
Predictor taxa with highest positive associations to eight focal algal lineages by CAP.

**Figure 3:**
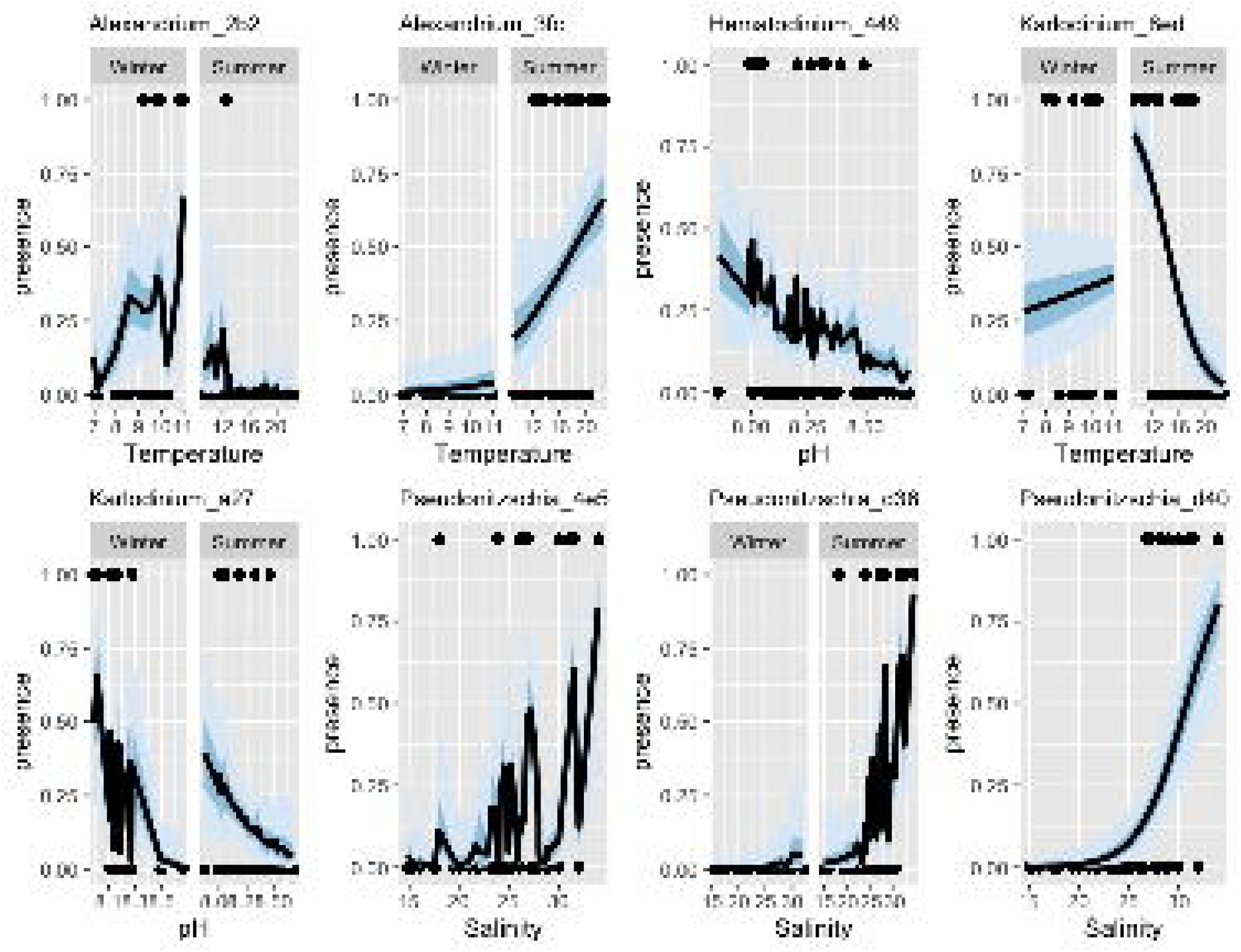
Best-fit logistic models for eight focal algal lineages. Here, probability of presence is shown as a function of the single most influential environmental variable in the model, along with overall model means (lines) and 50% and 95% credibility intervals (shaded areas). Where two-panel figures are shown for a taxon, the best-fit model included a slope term that varied by season.

### Biological Context

To identify the biological community associated with our focal lineages, we searched for co-occurring taxa using a canonical analysis of principal coordinates (CAP) (Anderson & Willis, 2003). Constraining this multivariate analysis according to the presence or absence of each potential harmful algal variant revealed no striking patterns of association across taxa (Tables S4-S11), but rather helped to identify individual community members particularly likely or unlikely to co-occur with our focal lineages. These associated community members were those with the strongest deviations from 0 on the CAP1 axis (Table 3), and included lineages of *Ditylum*, a centric diatom, *Prasinoderma*, a non-harmful green algae, *Saxidomus*, a clam, *Balanus*, a barnacle, and *Calanus*, a copepod.

For each of our focal lineages, adding the most closely associated predictor taxon improved model fit even after accounting for the additional model complexity (Table 4). Thus, including information about co-occurring organisms alongside baseline environmental covariates substantially increased our ability to predict the presence of these potential harmful algal taxa within the scope of our sampling. For example, Hematodinium_449 occurs somewhat stochastically in space and time (Figure 2) and is not strongly associated with environmental covariates (Table 2). However, the CAP analysis revealed that a haplotype from the clam genus *Saxidomus* (likely the species *S. giganteus* (butter clam), given the sampling location), was routinely found in samples in which *Hematodinium* also occurred (Table 3). Adding *Saxidomus* as a term in the previous best-fit model more than doubles the model’s overall accuracy (Figure 4). All else being equal, when Saxidomus DNA is detected, it more than quadruples the likelihood of *Hematodinium* being detected (0.08 vs. 0.45 mean detection probability), making this biological variable a far better predictor than the measured environmental variables alone.

**Table 4:** Terms of the best-fit models combining environmental variables and most closely associated biological taxon for eight focal algal lineages.

| Taxon | NA |
| --- | --- |
| Alexandrium_2b2 | Temperature + pH + (Intercept \| Ditylum presence) |
| Alexandrium_3fc | (Intercept + Temperature \| Season) + (Intercept \| Prasinoderma presence) |
| Hematodinium_449 | pH + Temperature + (Intercept \| Saxidomus presence) |
| Karlodinium_8ed | (Temperature \| Season) + (Intercept \| Ditylum presence) |
| Karlodinium_a27 | pH + (Salinity \| Season) + (Intercept \| Balanus presence) |
| Pseudonitzschia_4e5 | Salinity + Temperature + pH + (Intercept \| Calanus presence) |
| Pseudonitzschia_d36 | Salinity + pH + (Intercept \| Calanus presence) |
| Pseudonitzschia_d40 | Salinity + (Intercept \| Ditylum presence) |

**Figure 4:**
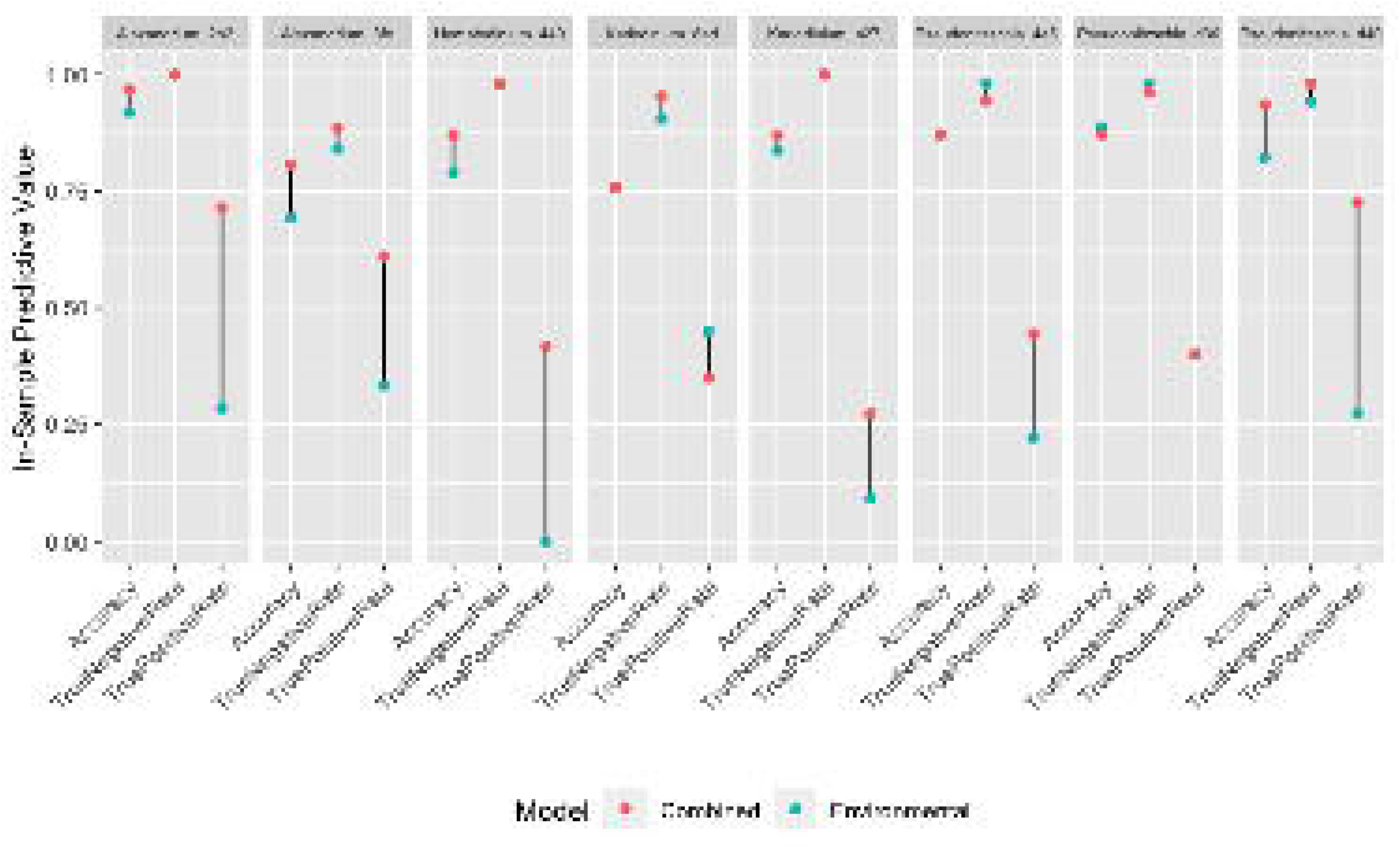
In-sample predictive value of best-fit models for eight focal algal lineages, as measured by accuracy, true-negative rate, and true-positive rate. Red dots indicate values for models combining environmental information with a single associated predictor taxon; blue dots indicate values for models with environmental information alone. Where only a single dot is visible, models produced equivalent results.

Performing the same analysis for each of our focal lineages yields similar results (Figure 4), demonstrating an overall model accuracy substantially above environmental covariates alone for most potential harmful algal variants. Specifically, adding these candidate indicator taxa improved the true-positive rate of detection for six of the eight models (Karlodinium_8ed and Pseudonitzschia_d36 are the exceptions), which accounted for the increase in overall accuracy across models.

## Discussion

Here, we use genetic monitoring to highlight a wide variety of putative harmful algal taxa from a larger set of several hundred taxa in intertidal and nearshore marine habitats. Within these prospective harmful algal groups, we find several variants from lineages that are unexpected in the study area (*Karlodinium*, *Hematodinium*), in addition to cryptic lineages of others (most notably, *Alexandrium* sp.), and multiple variants of economically important taxa (e.g. *Pseudo-nitzschia*). Our time-series sampling indicates different seasonal patterns and attendant associations of sea-surface temperature, pH, and salinity for some of these lineages, but on the whole, models using purely environmental covariates offer poor predictive value. We therefore use a constrained ordination to identify taxa (both algal and non-algal) that commonly occur in association with our focal lineages; adding only a single prospective biological indicator taxon greatly improved most of the predictive models.

### Detecting Expected and Unexpected Harmful Algal Taxa

eDNA metabarcoding has a number of distinct advantages relative to common techniques currently used to identify harmful algae. First, this technique can reveal a diversity of potential harmful algal variants present, rather than targeting specific species. In our survey of intertidal and nearshore communities using broad-spectrum eukaryotic mitochondrial COI primers, the taxa we identified were largely consistent with what we expected *a priori*, in that we found dozens of variants with excellent overlap from records of known local harmful algal genera (Table 1; Moestrup et al., 2020; Horner & Postel, 1993; Horner, Garrison & Plumley, 1997; Trainer et al., 2016), such as three from the genus *Pseudo-nitzschia*, which is represented by multiple species in the Puget Sound (Hubbard, Olson & Armbrust, 2014). While confirming expectations demonstrates the reliability of eDNA metabarcoding for identification, detecting unexpected taxa underscores the ability of this technique to reveal novel lineages, range-shifts, or nascent invasions with potentially profound ecological and economic consequences. For example, the genus *Alexandrium*, though known to cause paralytic shellfish poisoning in the region (Trainer et al., 2016), was not previously understood to have two distinct seasonal forms (see below). Additionally, members of *Karlodinium* produce karlotoxins responsible for fish kills in the United States and globally (e.g. *Karlodinium veneficum* (Place et al., 2012)), yet this genus is not reported from Puget Sound in the peer-reviewed scientific literature, despite having been noted by a local monitoring program (Amelia Kolb Gabriela Hannach & Swanson, 2016). Similarly, member(s) of the genus *Hematodinium*, which have caused massive losses for the tanner crab (*Chionoecetes bairdi*) and snow crab (*C. opilio*) fisheries in the United States (Meyers et al., 1987, 1996; Wood et al., 2017) and among other species worldwide (Stentiford & Shields, 2005), have also not previously been reported within Puget Sound in the peer-reviewed scientific literature.

Additionally, eDNA metabarcoding can help to standardize data collection and analysis across studies. Here we employ samples collected with two distinct methods by two groups: intertidal water was gathered on foot and by hand, whereas nearshore water was gathered by boat on the Washington Ocean Acidification Cruise in a more routinized process. Nonetheless, potential harmful algal taxa identified in nearshore surface samples were all found within the larger intertidal dataset as well, suggesting excellent agreement despite differences in methods and personnel. Such congruence between studies also arises from how the data are analyzed: eDNA sequences from multiple studies can consistently identify cryptic taxa by sequence (e.g. Uchii, Doi & Minamoto, 2016; Thomsen & Willerslev, 2015), and can undergo identical taxonomic analyses that are not subject to differences in interpretation via morphology (Proschold & Leliaert, 2007). However, we note that the success of eDNA studies rests heavily on the shoulders of expert taxonomists: without their contributions to the identification of specimens with sequences in databases, it is impossible to link a fragment of DNA found in the water to an organism (Manoylov, 2014; Zimmermann et al., 2014).

### Species Distributions in Space and Time

Our spatial and temporal data indicate that many putative harmful algal taxa are constitutively present in the intertidal and nearshore environment, or at the very least are routinely detectable (Table 1; Figure 2). Although challenges in relating sequence counts to absolute organism abundances limit the utility of eDNA metabarcoding for precise measurement of bloom intensity, the ability of eDNA to reveal harmful algal taxa even when cell counts are much lower than bloom conditions can be advantageous. For example, we detect *Alexandrium* variants year-round in Hood Canal, including winters, when recorded *Alexandrium* blooms are rare, but when fisheries closures due to presence of paralytic shellfish toxin do exist (Trainer et al., 2003; Moore, Mantua & Salathe Jr, 2011). When paired with more traditional methods, this tool therefore provides another layer of information regarding the behavior of harmful algae, and at the very least can indicate a temporal and spatial starting place for more time-, labor-, and taxonomic expertise-intensive counting strategies.

Sampling many taxa over time and space additionally facilitates important within- and cross-species comparisons (Figure 2). Here, such comparisons reveal two lineages of *Alexandrium* with different temporal patterns, and completely non-overlapping distributions. Although taxonomic revisions of the *Alexandrium tamarense* species complex – and competing classifications of local taxa (Lilly, Halanych & Anderson, 2007; John et al., 2014) – make it impossible to identify the variants present in our survey without additional information, amino acid differences in the COI sequence of the two lineages alongside temporal distribution information suggest that they may represent distinct species, rather than haplotypes. Regardless, recognizing two distinct *Alexandrium* lineages with opposite seasonal dynamics is likely to be important to local monitoring and research programs aiming to identify risk of saxitoxin poisoning (e.g. Amelia Kolb Gabriela Hannach & Swanson, 2016; Trainer et al., 2016). Previous work on *Alexandrium* within the Puget Sound found a lower limit for toxic bloom events at 13° C (Nishitani & Chew, 1984), with recent work identifying higher growth of cells above 17-18° C (Bill et al., 2016), and more frequent blooms accompanying warmer air and sea surface temperatures over multiple decades (Moore et al., 2009; Moore, Mantua & Salathe Jr, 2011). Alexandrium_2b2, present nearly exclusively in winter, cold-water samples, does not match the profile expected given these studies, suggesting either that it does not bloom frequently and/or that it is a recent introduction to the local algal community, whose role is not yet appreciated.

Our dataset also underscores the dynamism and diversity of harmful algal taxa in local waters (Figure 2). For example, some unexpected variants in our dataset (e.g. Hematodinium_449 and Karlodinium_a27) are highly variable in space and time, while others are more consistent and widespread (e.g Karlodinium 8ed, which appears in 20 of 63 samples). These results indicate our method may reveal transient appearances as well as the well-established presence of understudied taxa in the region. Notably, members of the genus *Hematodinium* have been reported on the west coast of North America in the Bering Sea and Southeast Alaska (Meyers et al., 1987, 1996; Jensen et al., 2010; Small, 2012). Likewise, *Karlodinium venificum* was first documented in San Francisco Bay in 2005, with continuing observations over the following decade (Nejad, Schraga & Cloern), but not elsewhere on the West Coast of North America (Moestrup et al., 2020). This study therefore adds evidence to reports of these taxa in nearby locales, suggesting that more work is necessary to track their presence in the Puget Sound region, where it is possible that they might impact important fisheries in the future.

Nucleotide variants from the genus *Pseudo-nitzschia* identified here are not unexpected; multiple species from this genus have been described locally using both visual (e.g. Trainer et al., 2016) and molecular tools (e.g. Hubbard, Olson & Armbrust, 2014). In the Western Pacific, clades of a single *Pseudo-nitzschia* species (*P. pungens*) with distinct ecological niches and hybrid zones have been documented (Kim et al., 2018); it is interesting to note that here we similarly find two *Pseudo-nitzschia* variants with distinct nucleotide but identical amino acid sequences at the COI locus. Based on their overlap in time and space, it is likely these represent haplotype lineages that have begun to diverge but are not yet distinct species. Additionally, the specific local antecedents of toxicity in *Pseudo-nitzschia* are still under study (e.g. Zhu et al., 2017; Trick et al., 2018), with production of domoic acid historically limited to the outer coast of Washington (Trainer et al., 2017), but recently moving into Puget Sound (Trainer et al., 2007). Revealing the diversity and pattern of genetic variants present in time and space by eDNA metabarcoding might thus support efforts to better characterize the causes underlying dangerous and costly *Pseudo-nitzschia* bloom events.

### Environmental and Biological Context

Quantitative models of each focal lineage with respect to environmental variables (Table 2), motivated by taxon-specific patterns in space and time (Figure 2), yield a synoptic view of the occurrence of many potentially harmful algal taxa in the region – a perspective that is otherwise not easily achievable, though many detailed quantitative models have been built for individual harmful algal species (e.g. Moore et al., 2015; Hubbard, Olson & Armbrust, 2014). Across taxa, we note that values expected with global climate change (higher sea surface temperature) and ocean acidification (lower pH) are typically associated with increases in the occurrence of our focal lineages, such as Alexandrium_2b2, Alexandrium_3fc, Hematodinium_449, and Karlodinium_a27. These results in sum align with other studies of local harmful algal taxa, suggesting future scenarios will involve greater seasonal windows of opportunity for toxic bloom events (e.g. Moore, Mantua & Salathe Jr, 2011; Trainer et al., 2020).

Overall, however, the quantitative models we have built with environmental variables alone have low accuracy, and universally fail to predict a majority of occurrences for our focal lineages (Table 2). The difficulty of predicting the presence of harmful algal taxa and their blooms based on a limited number of environmental covariates is not atypical; building accurate models of these species’ ecology has long been a challenge for the field (Flynn & McGillicuddy, 2018), even when many more environmental covariates are considered. In this study, the variant Hematodinium_449 is an extreme representative of this challenge; our environmental model does not predict its presence correctly even once (Table 2), though it appears in 12 of 63 samples gathered. Similarly, Karlodinium_a27 and Pseudonitzchia_4e5 models both have true positive rates less than 0.25, according to their respective environmental models (Table 2). These models are consequently worse than uniformative; they can be actively misleading by inaccurately predicting the absence of harmful species.

Fortunately, eDNA metabarcoding provides an additional layer of information regarding the context of harmful algal taxa: the surrounding biological community members (both algal and non-algal). Specifically, the choice of universal eukaryotic primers amplifying a common molecular marker (mitochondrial COI) allows us to identify over 600 taxa in total, from more than 250 genera. These data enable us to associate the presence and absence of individual focal variants with a wide range of eukaryotes by CAP (Tables 3 and S4-S11). Studies such as this one that examine harmful algae within the breadth of their biological communities are rare, but provide an opportunity to improve prediction (e.g. Banerji et al., 2019). Here, we see that adding only the single best predictor taxon to our quantitative models of focal lineages universally improves their fit (Table 5), justifying added model complexity. Although it may appear circular to identify co-occurring species in the dataset and subsequently add them into the model of the same dataset, use of WAIC allows us to assess the value of additional information for future out-of-sample data, generating testable hypotheses for harmful algal indicator species. As an example, adding the presence of an easily surveyed indicator taxon, a *Saxidomus* clam, improves the prediction of Hematodinium_449 dramatically, in particular the true-positive measure essential to management (Figure 4). The majority of focal lineages examined here similarly show an improvement in model accuracy driven by large increases in the true positive rate with addition of biological information.

## Conclusion

In this study, we employ COI universal primers (Leray et al., 2013) to simultaneously identify dozens of potentially harmful algal variants, as well as hundreds of other local taxa comprising the biological context for these harmful algae. The broad nature of eDNA metabarcoding surveys allows us to track both expected and unexpected taxa, and the distribution in time and space of eight focal variants from the genera *Alexandrium*, *Hematodinium*, *Karlodinium*, and *Pseudo-nitzschia* suggests the constitutive presence of harmful algae in the study region, as well the possibility of nascent range shifts, invasions, and/or ongoing evolutionary divergence. Building individual quantitative models for each of eight focal lineages, we find that many variants are likely to become more common under conditions of higher sea surface temperature and ocean acidification, but note that models using environmental covariates alone have low explanatory power. Adding even a single associated member of the biological community, however, improves most models, and in particular boosts the true positive rates useful for prediction of harmful algal taxa in the field. eDNA metabarcoding is hence an opportunity to reveal harmful algae outside of bloom events and expected ranges, to map phylogenetic complexity underlying HAB dynamics, to interrogate the relevant environmental context in an era of global change, and to improve models of harmful algal prediction with inclusion of the biological milieu.

## Acknowledgements

We thank the Washington Ocean Acidification Center (WOAC) National Oceanic and Atmospheric Administration Harmful Algal Blooms (NOAA HABs) program cruise, and in particular Simone Alin for her provision of samples and environmental data, as well as Terrie Klinger and Jan Newton for their expert comments on the manuscript. We thank our colleagues at the NOAA Northwest Fisheries Science Center, including the Conservation Biology Molecular Genetics Laboratory for their use of the Illumina MiSeq, as well as Linda Rhodes, Stephanie Moore, Vera Trainer, and Nic Adams for their direction of the analysis in its early stages. We thank Micah Horwith and the Washington State Department of Natural Resources for field assistance. Finally, we thank Alex Gagnon, in whose lab we performed carbonate chemistry analysis for the intertidal dataset, and the lab groups in the University of Washington’s Center for Environmental Genomics, with whom we share a work space.

## References

Ahn Y-H., Shanmugam P., Ryu J-H., Jeong J-C. 2006. Satellite detection of harmful algal bloom occurrences in Korean waters. Harmful Algae 5:213–231.

Alin SR., Newton J., Greeley D., Curry B., Herndon J., Ostendorf ML., Mitchell-Morton A., Feely RA. 2019a. Dissolved inorganic carbon (DIC), total alkalinity (TA), temperature, salinity, oxygen, nutrient, and CTD data collected from discrete profile measurements during Puget Sound cruise CAB1079 (EXPOCODE 33CB20170911) on R/V Clifford A. Barnes from 2017-09-11 to 2017-09-15 (NCEI Accession 0206674). DIC, TA, temperature, salinity, and CTD data from sites P12 and P402 at depth = 0. NOAA National Centers for Environmental Information. Dataset. DOI: https://doi.org/10.25921/nhgn-7685.

Alin SR., Newton J., Greeley D., Curry B., Herndon J., Ostendorf ML., Mitchell-Morton A., Feely RA. 2019b. Dissolved inorganic carbon (DIC), total alkalinity (TA), temperature, salinity, oxygen, nutrient, and CTD data collected from discrete profile measurements during Puget Sound cruise RC001 (EXPOCODE 33IY20180407) on R/V Rachel Carson from 2018-04-07 to 2018-04-11 (NCEI Accession 0206802). DIC, TA, temperature, salinity, and CTD data from sites P8, P12, and P402 at depth = 0. NOAA National Centers for Environmental Information. Dataset. DOI: https://doi.org/10.25921/nk76-he39.

Alin SR., Newton J., Greeley D., Curry B., Herndon J., Ostendorf ML., Mitchell-Morton A., Feely RA. 2019c. Dissolved inorganic carbon (DIC), total alkalinity (TA), temperature, salinity, oxygen, nutrient, and CTD data collected from discrete profile measurements during Puget Sound cruise RC007 (EXPOCODE 33IY20180911) on R/V Rachel Carson from 2018-09-11 to 2018-09-15 (NCEI Accession 0206804). DIC, TA, temperature, salinity, and CTD data from sites P11, P12, and P14 at depth = 0. NOAA National Centers for Environmental Information. Dataset. DOI: https://doi.org/10.25921/xp63-pm64.

Al-Tebrineh J., Mihali TK., Pomati F., Neilan BA. 2010. Detection of saxitoxin-producing cyanobacteria and Anabaena circinalis in environmental water blooms by quantitative PCR. Appl. Environ. Microbiol. 76:7836–7842.

Amelia Kolb Gabriela Hannach., Swanson L. 2016. Marine Phytoplankton Monitoring Program Sampling and Analysis Plan. Seattle, WA: King County Department of Natural Resources; Parks.

Anderson MJ., Willis TJ. 2003. Canonical analysis of principal coordinates: a useful method of constrained ordination for ecology. Ecology 84:511–525.

Antonella P., Luca G. 2013. The quantitative real-time PCR applications in the monitoring of marine harmful algal bloom (HAB) species. Environmental Science and Pollution Research 20:6851–6862.

Banerji A., Bagley MJ., Shoemaker JA., Tettenhorst DR., Nietch CT., Allen HJ., Santo Domingo JW. 2019. Evaluating putative ecological drivers of microcystin spatiotemporal dynamics using metabarcoding and environmental data. Harmful Algae 86:84–95.

Bill BD., Moore SK., Hay LR., Anderson DM., Trainer VL. 2016. Effects of temperature and salinity on the growth of *Alexandrium* (Dinophyceae) isolates from the Salish Sea. Journal of phycology 52:230–238.

Busch DS., Harvey CJ., McElhany P. 2013. Potential impacts of ocean acidification on the Puget Sound food web. ICES Journal of Marine Science 70:823–833. DOI: 10.2307/4451315.

Buskey EJ. 2008. How does eutrophication affect the role of grazers in harmful algal bloom dynamics? Harmful Algae 8:152–157.

Callahan BJ., McMurdie PJ., Rosen MJ., Han AW., Johnson AJA., Holmes SP. 2016. DADA2: high-resolution sample inference from Illumina amplicon data. Nature methods 13:581.

Campbell L., Olson RJ., Sosik HM., Abraham A., Henrichs DW., Hyatt CJ., Buskey EJ. 2010. First Harmful *Dinophysis* (Dinophyceae, Dinophysales) Bloom in the U.S. is Revealed by Automated Imaging Flow Cytometry. Journal of Phycology 46:66–75.

Cembella AD., Lewis NI., Quilliam MA. 2000. The marine dinoflagellate *Alexandrium ostenfeldii* (Dinophyceae) as the causative organism of spirolide shellfish toxins. Phycologia 39:67–74. DOI: 10.2216/i0031-8884-39-1-67.1.

Cho K., Kasaoka T., Ueno M., Basti L., Yamasaki Y., Kim D., Oda T. 2017. Haemolytic activity and reactive oxygen species production of four harmful algal bloom species. European Journal of Phycology 52:311–319.

Curd EE., Gold Z., Kandlikar GS., Gomer J., Ogden M., O’Connell T., Pipes L., Schweizer TM., Rabichow L., Lin M., Others. 2019. Anacapa Toolkit: an environmental DNA toolkit for processing multilocus metabarcode datasets. Methods in Ecology and Evolution 10:1469–1475.

Deiner K., Bik HM., Mächler E., Seymour M., Lacoursière-Roussel A., Altermatt F., Creer S., Bista I., Lodge DM., De Vere N., Others. 2017. Environmental DNA metabarcoding: Transforming how we survey animal and plant communities. Molecular ecology 26:5872–5895.

Dickson AG., Sabine CL., Christian JR. 2007. Guide to best practices for ocean co2 measurements. North Pacific Marine Science Organization.

Dìaz PA., Álvarez A., Varela D., Pérez-Santos I., Dìaz M., Molinet C., Seguel M., Aguilera-Belmonte A., Guzmán L., Uribe E., Others. 2019. Impacts of harmful algal blooms on the aquaculture industry: Chile as a case study. Perspectives in Phycology.

Erdner DL., Percy L., Keafer B., Lewis J., Anderson DM. 2010. A quantitative real-time PCR assay for the identification and enumeration of Alexandrium cysts in marine sediments. Deep Sea Research Part II: Topical Studies in Oceanography 57:279–287.

Feely RA., Alin SR., Newton J., Sabine CL., Warner M., Devol A., Krembs C., Maloy C. 2010. The combined effects of ocean acidification, mixing, and respiration on pH and carbonate saturation in an urbanized estuary. Estuarine, Coastal and Shelf Science 88:442–449. DOI: 10.1016/j.ecss.2010.05.004.

Ferrante M., Conti GO., Fiore M., Rapisarda V., Ledda C. 2013. Harmful Algal Blooms in the Mediterranean Sea: Effects on Human Health. EuroMediterranean Biomedical Journal 8.

Field CB., Barros VR., Mastrandrea MD., Mach KJ., Abdrabo MA-K., Adger WN., Anokhin YA., Anisimov OA., Arent DJ., Barnett J., Others. 2014. Summary for Policy Makers: Working group 11 contribution to the fifth assessment report of the Intergovernmental panel on Climate Change.

Flynn KJ., McGillicuddy DJ. 2018. Modeling marine harmful algal blooms: Current status and future prospects. Harmful Algal Blooms: A Compendium Desk Reference:115–134.

Fu FX., Tatters AO., Hutchins DA. 2012. Global change and the future of harmful algal blooms in the ocean. Marine Ecology Progress Series 470:207–233.

Gallego R. demultiplexer_for_DADA2.

Gasteiger E., Gattiker A., Hoogland C., Ivanyi I., Appel RD., Bairoch A. 2003. ExPASy: The proteomics server for in-depth protein knowledge and analysis. Nucleic acids research 31:3784–3788.

Gattuso J-P., Magnan A., Billé R., Cheung WWL., Howes EL., Joos F., Allemand D., Bopp L., Cooley SR., Eakin CM., Others. 2015a. Contrasting futures for ocean and society from different anthropogenic CO2 emissions scenarios. Science 349:aac4722.

Gattuso J-P., Epitalon J-M., Lavigne H., Orr J., Gentili B., Hofmann A., Proye A., Soetaert K., Rae J. 2015b. seacarb: Seawater carbonate chemistry. R package version 3.

Gobler CJ., Doherty OM., Hattenrath-Lehmann TK., Griffith AW., Kang Y., Litaker RW. 2017. Ocean warming since 1982 has expanded the niche of toxic algal blooms in the North Atlantic and North Pacific oceans. Proceedings of the National Academy of Sciences 114:4975–4980.

Gobler CJ. 2020. Climate change and harmful algal blooms: insights and perspective. Harmful Algae 91:101731.

Graneli E., Lipiatou E. 2002. EUROHAB, Part B, Research and infrastructural needs, National European and international programmes.

Grattan LM., Holobaugh S., Morris Jr JG. 2016. Harmful algal blooms and public health. Harmful algae 57:2–8.

Griffith AW., Gobler CJ. 2020. Harmful algal blooms: a climate change co-stressor in marine and freshwater ecosystems. Harmful Algae 91:101590.

Grzebyk D., Audic S., Lasserre B., Abadie E., Vargas C de., Bec B. 2017. Insights into the harmful algal flora in northwestern Mediterranean coastal lagoons revealed by pyrosequencing metabarcodes of the 28S rRNA gene. Harmful algae 68:1–16.

Horner RA., Garrison DL., Plumley FG. 1997. Harmful algal blooms and red tide problems on the US west coast. Limnology and Oceanography 42:1076–1088.

Horner RA., Postel JR. 1993. Toxic diatoms in western Washington waters (US west coast). In: Twelfth international diatom symposium. Springer, 197–205.

Hubbard KA., Olson CE., Armbrust EV. 2014. Molecular characterization of *Pseudo-nitzschia* community structure and species ecology in a hydrographically complex estuarine system (Puget Sound, Washington, USA). Marine ecology progress series 507:39–55.

Jensen PC., Califf K., Lowe V., Hauser L., Morado JF. 2010. Molecular detection of *Hematodinium* sp. in Northeast Pacific *Chionoecetes* spp. and evidence of two species in the Northern Hemisphere. Diseases of aquatic organisms 89:155–166.

John U., Litaker RW., Montresor M., Murray S., Brosnahan ML., Anderson DM. 2014. Formal revision of the textitAlexandrium tamarense species complex (Dinophyceae) taxonomy: the introduction of five species with emphasis on molecular-based (rDNA) classification. Protist 165:779–804.

Kelly RP., Gallego R., Jacobs-Palmer E. 2018. The effect of tides on nearshore environmental DNA. PeerJ. 2018. DOI: 10.7717/peerj.4521.

Kelly RP., Shelton AO., Gallego R. 2019. Understanding PCR processes to draw meaningful conclusions from environmental DNA studies. Scientific reports 9:1–14.

Khan S., Arakawa O., Onoue Y. 1997. Neurotoxins in a toxic red tide of *Heterosigma akashiwo* (Raphidophyceae) in Kagoshima Bay, Japan. Aquaculture Research 28:9–14.

Kim JH., Wang P., Park BS., Kim J-H., Patidar SK., Han M-S. 2018. Revealing the distinct habitat ranges and hybrid zone of genetic sub-populations within *Pseudo-nitzschia pungens* (Bacillariophyceae) in the West Pacific area. Harmful algae 73:72–83.

Kotaki Y., Furio EF., Bajarias FA., Satake M., Lundholm N., Koike K., Sato S., Fukuyo Y., Kodama M. 2006. New stage of the study on domoic acid-producing diatoms–a finding of *Nitzschia navis-varingica* that produces domoic acid derivatives as major toxin components. Coastal Marine Science 30:116–120.

Lahoz-Monfort JJ., Guillera-Arroita G., Tingley R. 2016. Statistical approaches to account for false-positive errors in environmental dna samples. Molecular Ecology Resources 16:673–685.

Lapworth C., Hallegraeff GMJ., Ajani PA. 2001. Identification of domoic-acid-producing *Pseudo-nitzschia* species in Australian waters. Harmful Algal Blooms 2000 Proceedings of the Ninth International Conference on Harmful Algal Blooms:38–41.

Leray M., Yang JY., Meyer CP., Mills SC., Agudelo N., Ranwez V., Boehm JT., Machida RJ. 2013. A new versatile primer set targeting a short fragment of the mitochondrial COI region for metabarcoding metazoan diversity: application for characterizing coral reef fish gut contents. Frontiers in zoology 10:34.

Lewitus AJ., Horner RA., Caron DA., Garcia-Mendoza E., Hickey BM., Hunter M., Huppert DD., Kudela RM., Langlois GW., Largier JL., Others. 2012. Harmful algal blooms along the North American west coast region: History, trends, causes, and impacts. Harmful algae 19:133–159.

Lilly EL., Halanych KM., Anderson DM. 2007. Species boundaries and global biogeography of the Alexandrium tamarense complex (Dinophyceae) 1. Journal of Phycology 43:1329–1338.

Lindberg K., Moestrup Ø., Daugbjerg N. 2005. Studies on woloszynskioid dinoflagellates i: *Woloszynskia coronata* re-examined using light and electron microscopy and partial lsu rDNA sequences, with description of *tovellia* gen. Nov. And *jadwigia* gen. Nov.(Tovelliaceae fam. Nov.). Phycologia 44:416–440.

Lopez CB., Jewett EB., Dortch Q., Walton BT., Hudnell HK. 2008. Scientific assessment of freshwater harmful algal blooms.

Loureiro S., Reñé A., Garcés E., Camp J., Vaqué D. 2011. Harmful algal blooms (HABs), dissolved organic matter (DOM), and planktonic microbial community dynamics at a near-shore and a harbour station influenced by upwelling (SW Iberian Peninsula). Journal of Sea Research 65:401–413.

Mahé F., Rognes T., Quince C., Vargas C de., Dunthorn M. 2015. Swarm v2: highly-scalable and high-resolution amplicon clustering. PeerJ 3:e1420.

Manoylov KM. 2014. Taxonomic identification of algae (morphological and molecular): Species concepts, methodologies, and their implications for ecological bioassessment. Journal of phycology 50:409–424.

Martin M. 2011. Cutadapt removes adapter sequences from high-throughput sequencing reads. EMBnet.journal 17:pp—–10.

Mauger G., Casola J., Morgan H., Strauch R., Jones B., Curry B., Busch Isaksen T., Whitely Binder L., Krosby M., Snover A. 2015. State of knowledge: Climate change in Puget Sound. NOAA and the University of Washington Climate Impacts Group.

McElreath R. 2020. Statistical rethinking: A Bayesian course with examples in R and Stan. CRC press.

Meyers TR., Koeneman TM., Botelho C., Short S., others. 1987. Bitter crab disease: a fatal dinoflagellate infection and marketing problem for Alaskan Tanner crabs *Chionoecetes bairdi*. Diseases of Aquatic Organisms 3:195–216.

Meyers T., Morado J., Sparks A., Bishop G., Pearson T., Urban D., Jackson D. 1996. Distribution of bitter crab syndrome in Tanner crabs (*Chionoecetes bairdi, C. opilio*) from the Gulf of Alaska and the Bering Sea. Diseases of aquatic organisms 26:221–227.

Moestrup Ø., Akselmann-Cardella R., Churro C., Fraga S., Hoppenrath M., Iwataki M., Larsen J., Lundholm N., Zingone A. 2020. IOC-UNESCO Taxonomic Reference List of Harmful Micro Algae. DOI: doi:10.14284/362.

Moore SK., Mantua NJ., Hickey BM., Trainer VL. 2009. Recent trends in paralytic shellfish toxins in Puget Sound, relationships to climate, and capacity for prediction of toxic events. Harmful Algae 8:463–477.

Moore SK., Johnstone JA., Banas NS., Salathe Jr EP. 2015. Present-day and future climate pathways affecting Alexandrium blooms in Puget Sound, WA, USA. Harmful Algae 48:1–11.

Moore SK., Cline MR., Blair K., Klinger T., Varney A., Norman K. 2019. An index of fisheries closures due to harmful algal blooms and a framework for identifying vulnerable fishing communities on the US West Coast. Marine Policy 110:103543.

Moore SK., Mantua NJ., Salathe Jr EP. 2011. Past trends and future scenarios for environmental conditions favoring the accumulation of paralytic shellfish toxins in puget sound shellfish. Harmful Algae 10:521–529.

Murray SA., Wiese M., Stüken A., Brett S., Kellmann R., Hallegraeff G., Neilan BA. 2011. sxtA-based quantitative molecular assay to identify saxitoxin-producing harmful algal blooms in marine waters. Applied and Environmental Microbiology 77:7050–7057.

National Centers for Coastal Ocean Science. 2014. Working with state to document first occurrence of harmful karenia mikimotoi algae in Alaskan waters.

Nejad ES., Schraga TS., Cloern JE. Phytoplankton Species Composition, Abundance and Cell Size in San Francisco Bay: Microscopic Analyses of USGS Samples, beginning in 2014 (ver. 2.0, April 2019). DOI: https://doi.org/10.5066/F7C24VB2.

Nishitani L., Chew KK. 1984. Recent developments in paralytic shellfish poisoning research. Aquaculture 39:317–329.

O’Donnell JL., Kelly RP., Lowell NC., Port JA. 2016. Indexed PCR primers induce template-specific bias in large-scale DNA sequencing studies. PloS one 11:e0148698.

Oksanen J., Blanchet FG., Kindt R., Legendre P., Minchin PR., O’hara RB., Simpson GL., Solymos P., Stevens MHH., Wagner H., Others. 2013. Package “vegan”. Community ecology package, version 2.

Peperzak L., Poelman M. 2008. Mass mussel mortality in the netherlands after a bloom of phaeocystis globosa (prymnesiophyceae). Journal of Sea Research 60:220–222.

Pierce RH., Kirkpatrick GJ. 2001. Innovative techniques for harmful algal toxin analysis. Environmental Toxicology and Chemistry: An International Journal 20:107–114.

Place AR., Bowers HA., Bachvaroff TR., Adolf JE., Deeds JR., Sheng J. 2012. *Karlodinium veneficum*—The little dinoflagellate with a big bite. Harmful Algae 14:179–195.

Plummer M., Stukalov A., Denwood M., Plummer MM. 2016. Package ‘rjags’. Vienna, Austria.

Proschold T., Leliaert F. 2007. Systematics of the green algae: Conflict of classic and modern approaches. Systematics Association Special Volume 75:123.

Renshaw MA., Olds BP., Jerde CL., McVeigh MM., Lodge DM. 2015. The room temperature preservation of filtered environmental DNA samples and assimilation into a phenol–chloroform–isoamyl alcohol DNA extraction. Molecular ecology resources 15:168–176.

Royle JA., Link WA. 2006. Generalized Site Occupancy Models Allowing for False Positive and False Negative Errors. Ecology 87:835–841.

Ruvindy R., Bolch CJ., MacKenzie L., Smith KF., Murray SA. 2018. qPCR assays for the detection and quantification of multiple paralytic shellfish toxin-producing species of *Alexandrium*. Frontiers in microbiology 9:3153.

Satake M., MacKenzie L., Yasumoto T. 1998. Identification of *Protoceratium reticulatum* as the biogenetic origin of yessotoxin. Natural Toxins 5:164–167.

Schnell IB., Bohmann K., Gilbert MTP. 2015. Tag jumps illuminated–reducing sequence-to-sample misidentifications in metabarcoding studies. Molecular Ecology Resources.

Shimizu Y., Alam M., Oshima Y., Fallon WE. 1975. Presence of four toxins in red tide infested clams and cultured *Gonyaulax tamarensis* cells. Biochemical and Biophysical Research Communications 66:731–737. DOI: 10.1016/0006-291X(75)90571-9.

Sievers F., Higgins DG. 2014. Clustal Omega, accurate alignment of very large numbers of sequences. In: Multiple sequence alignment methods. Springer, 105–116.

Simonsen S., Moestrup Ø. 1997. Toxicity tests in eight species of *Chrysochromulina* (Haptophyta). Canadian journal of botany 75:129–136.

Skjelbred B., Horsberg TE., Tollefsen KE., Andersen T., Edvardsen B. 2011. Toxicity of the ichthyotoxic marine flagellate *Pseudochattonella* (Dictyochophyceae, Heterokonta) assessed by six bioassays. Harmful Algae 10:144–154.

Small HJ. 2012. Advances in our understanding of the global diversity and distribution of *hematodinium* spp.–Significant pathogens of commercially exploited crustaceans. Journal of invertebrate pathology 110:234–246.

Stentiford GD., Shields JD. 2005. A review of the parasitic dinoflagellates *Hematodinium* species and *Hematodinium*-like infections in marine crustaceans. Diseases of aquatic organisms 66:47–70.

Taberlet P., Bonin A., Coissac E., Zinger L. 2018. Environmental DNA: For biodiversity research and monitoring. Oxford University Press.

Thomsen PF., Willerslev E. 2015. Environmental DNA–An emerging tool in conservation for monitoring past and present biodiversity. Biological conservation 183:4–18.

Tomlinson MC., Stumpf RP., Ransibrahmanakul V., Truby EW., Kirkpatrick GJ., Pederson BA., Vargo GA., Heil CA. 2004. Evaluation of the use of SeaWiFS imagery for detecting *Karenia brevis* harmful algal blooms in the eastern Gulf of Mexico. Remote Sensing of Environment 91:293–303.

Trainer VL. 2002. Harmful algal blooms on the US west coast. Harmful algal blooms in the PICES region of the North Pacific:89–118.

Trainer VL., Eberhart B-TL., Wekell JC., Adams NG., Hanson L., Cox F., Dowell J. 2003. Paralytic shellfish toxins in Puget Sound, Washington state. Journal of Shellfish Research 22:213–223.

Trainer VL., Cochlan WP., Erickson A., Bill BD., Cox FH., Borchert JA., Lefebvre KA. 2007. Recent domoic acid closures of shellfish harvest areas in washington state inland waterways. Harmful Algae 6:449–459.

Trainer VL., Wells ML., Cochlan WP., Trick CG., Bill BD., Batgh KA., Beall BF., Herndon J., Lundholm N. 2009. An ecological study of a massive bloom of toxigenic *Pseudo-nitzschia* cuspidata off the Washington State coast. Limnology and Oceanography 54:1461–1474. DOI: 10.4319/lo.2009.54.5.1461.

Trainer V., Moore L., Bill B., Adams N., Harrington N., Borchert J., Silva D da., Eberhart B-T. 2013. Diarrhetic shellfish toxins and other lipophilic toxins of human health concern in Washington State. Marine Drugs 11:1815–1835.

Trainer V., King T., Bill B., Runyan J. 2016. SoundToxins manual: Puget Sound Harmful Algal Bloom Monitoring Program.[Revised December 2015].

Trainer VL., Adams NG., Bill BD., Ayres DL., Forster ZR., Odell A., Eberhart B-T., Haigh N. 2017. *Pseudo-nitzschia* blooms in the northeastern Pacific Ocean. PICES Sci. Rep:37–48.

Trainer VL., Moore SK., Hallegraeff G., Kudela RM., Clement A., Mardones JI., Cochlan WP. 2020. Pelagic harmful algal blooms and climate change: Lessons from nature’s experiments with extremes. Harmful algae 91:101591.

Trainer VL., Yoshida T. 2014. Proceedings of the Workshop on Economic Impacts of Harmful Algal Blooms on Fisheries and Aquaculture. North Pacific Marine Science Organization (PICES).

Trick CG., Trainer VL., Cochlan WP., Wells ML., Beall B. 2018. The successional formation and release of domoic acid in a *Pseudo-nitzschia* bloom in the Juan de Fuca Eddy: A drifter study. Harmful algae 79:105–114.

Uchii K., Doi H., Minamoto T. 2016. A novel environmental dna approach to quantify the cryptic invasion of non-native genotypes. Molecular Ecology Resources 16:415–422.

Watanabe S. 2010. Asymptotic equivalence of Bayes cross validation and widely applicable information criterion in singular learning theory. Journal of Machine Learning Research 11:3571–3594.

Wells ML., Trainer VL., Smayda TJ., Karlson BSO., Trick CG., Kudela RM., Ishikawa A., Bernard S., Wulff A., Anderson DM., Others. 2015. Harmful algal blooms and climate change: Learning from the past and present to forecast the future. Harmful algae 49:68–93.

Wood K., Stratman JP., Messmer A., Palof KJ. 2017. Annual management report for the 2016/2017 southeast alaska and yakutat tanner crab fisheries. Alaska Department of Fish; Game, Divisions of Sport Fish; Commercial Fisheries.

Yang ZB., Takayama H., Matsuoka K., Hodgkiss IJ. 2000. *Karenia digitata* sp. nov.(Gymnodiniales, Dinophyceae), a new harmful algal bloom species from the coastal waters of west Japan and Hong Kong. Phycologia 39:463–470.

Yang CZ., Albright LJ. 1994. The harmful phytoplankter *Chaetoceros concavicornis* causes high mortalities and leucopenia in chinook salmon (*Oncorhynchus tshawytscha*) and coho salmon (*O. kisutch*). Canadian Journal of Fisheries and Aquatic Sciences 51:2493–2500.

Zhu Z., Qu P., Fu F., Tennenbaum N., Tatters AO., Hutchins DA. 2017. Understanding the blob bloom: Warming increases toxicity and abundance of the harmful bloom diatom *Pseudo-nitzschia* in California coastal waters. Harmful Algae 67:36–43.

Zimmermann J., Abarca N., Enk N., Skibbe O., Kusber W-H., Jahn R. 2014. Taxonomic reference libraries for environmental barcoding: A best practice example from diatom research. PloS one 9:e108793.

