## Supplemental Tables S1-S11 for "Environmental DNA Metabarcoding for Simultaneous Monitoring and Ecological Assessment of Many Harmful Algae"

Emily Jacobs-Palmer

Ramón Gallego

Kelly Cribari

Abigail Keller

Ryan P. Kelly

Table S1: Sample location information. For each site, abbreviation, location type (nearshore/intertidal), and latitude/longitude are given.

| Site | Abbreviation | Date | Location | Latitude | Longitude |
| --- | --- | --- | --- | --- | --- |
| Lilliwaup Tidelands State Park | LL | 201703 | Intertidal | 47.4614 | -123.1062 |
| Potlatch State Park | PO | 201703 | Intertidal | 47.3605 | -123.1562 |
| Salisbury Point County Park | SA | 201703 | Intertidal | 47.8565 | -122.606 |
| Triton Cove State Park | TR | 201703 | Intertidal | 47.608 | -122.9856 |
| Twanoh State Park | TW | 201703 | Intertidal | 47.3785 | -122.9749 |
| Lilliwaup Tidelands State Park | LL | 201705 | Intertidal | 47.4615 | -123.1062 |
| Potlatch State Park | PO | 201705 | Intertidal | 47.3605 | -123.1562 |
| Salisbury Point County Park | SA | 201705 | Intertidal | 47.8565 | -122.606 |
| Triton Cove State Park | TR | 201705 | Intertidal | 47.608 | -122.9856 |
| Twanoh State Park | TW | 201705 | Intertidal | 47.3785 | -122.9749 |
| Lilliwaup Tidelands State Park | LL | 201706 | Intertidal | 47.4615 | -123.1062 |
| Potlatch State Park | PO | 201706 | Intertidal | 47.3605 | -123.1562 |
| Salisbury Point County Park | SA | 201706 | Intertidal | 47.8565 | -122.606 |
| Triton Cove State Park | TR | 201706 | Intertidal | 47.608 | -122.9856 |
| Twanoh State Park | TW | 201706 | Intertidal | 47.3785 | -122.9749 |
| Lilliwaup Tidelands State Park | LL | 201707 | Intertidal | 47.4615 | -123.1062 |
| Potlatch State Park | PO | 201707 | Intertidal | 47.3605 | -123.1562 |
| Salisbury Point County Park | SA | 201707 | Intertidal | 47.8565 | -122.606 |
| Triton Cove State Park | TR | 201707 | Intertidal | 47.608 | -122.9856 |
| Twanoh State Park | TW | 201707 | Intertidal | 47.3785 | -122.9749 |
| Lilliwaup Tidelands State Park | LL | 201708 | Intertidal | 47.4615 | -123.1062 |
| Potlatch State Park | PO | 201708 | Intertidal | 47.3605 | -123.1562 |
| Salisbury Point County Park | SA | 201708 | Intertidal | 47.8565 | -122.606 |
| Triton Cove State Park | TR | 201708 | Intertidal | 47.608 | -122.9856 |
| Twanoh State Park | TW | 201708 | Intertidal | 47.3785 | -122.9749 |
| Lilliwaup Tidelands State Park | LL | 201709 | Intertidal | 47.4615 | -123.1062 |
| Potlatch State Park | PO | 201709 | Intertidal | 47.3605 | -123.1562 |
| Salisbury Point County Park | SA | 201709 | Intertidal | 47.8565 | -122.606 |
| Triton Cove State Park | TR | 201709 | Intertidal | 47.608 | -122.9856 |
| Twanoh State Park | TW | 201709 | Intertidal | 47.3785 | -122.9749 |
| P12 | NA | 201709 | Nearshore (WOAC) | 47.4253 | -123.108 |
| P402 | NA | 201709 | Nearshore (WOAC) | 47.3563 | -123.0237 |
| Lilliwaup Tidelands State Park | LL | 201711 | Intertidal | 47.4615 | -123.1062 |
| Potlatch State Park | PO | 201711 | Intertidal | 47.3605 | -123.1562 |
| Salisbury Point County Park | SA | 201711 | Intertidal | 47.8565 | -122.606 |
| Triton Cove State Park | TR | 201711 | Intertidal | 47.608 | -122.9856 |
| Twanoh State Park | TW | 201711 | Intertidal | 47.3785 | -122.9749 |
| Lilliwaup Tidelands State Park | LL | 201801 | Intertidal | 47.4615 | -123.1062 |
| Potlatch State Park | PO | 201801 | Intertidal | 47.3605 | -123.1562 |
| Salisbury Point County Park | SA | 201801 | Intertidal | 47.8565 | -122.606 |
| Triton Cove State Park | TR | 201801 | Intertidal | 47.608 | -122.9856 |
| Twanoh State Park | TW | 201801 | Intertidal | 47.3785 | -122.9749 |
| Lilliwaup Tidelands State Park | LL | 201803 | Intertidal | 47.4615 | -123.1062 |
| Potlatch State Park | PO | 201803 | Intertidal | 47.3605 | -123.1562 |
| Salisbury Point County Park | SA | 201803 | Intertidal | 47.8565 | -122.606 |
| Triton Cove State Park | TR | 201803 | Intertidal | 47.608 | -122.9856 |
| Twanoh State Park | TW | 201803 | Intertidal | 47.3785 | -122.9749 |
| P8 | NA | 201804 | Nearshore (WOAC) | 47.8975 | -122.6053 |
| P12 | NA | 201804 | Nearshore (WOAC) | 47.4253 | -123.108 |
| P402 | NA | 201804 | Nearshore (WOAC) | 47.3563 | -123.0237 |
| Lilliwaup Tidelands State Park | LL | 201805 | Intertidal | 47.4615 | -123.1062 |
| Potlatch State Park | PO | 201805 | Intertidal | 47.3605 | -123.1562 |
| Salisbury Point County Park | SA | 201805 | Intertidal | 47.8565 | -122.606 |
| Triton Cove State Park | TR | 201805 | Intertidal | 47.608 | -122.9856 |
| Twanoh State Park | TW | 201805 | Intertidal | 47.3785 | -122.9749 |
| Lilliwaup Tidelands State Park | LL | 201807 | Intertidal | 47.4615 | -123.1062 |
| Potlatch State Park | PO | 201807 | Intertidal | 47.3605 | -123.1562 |
| Salisbury Point County Park | SA | 201807 | Intertidal | 47.8565 | -122.606 |
| Triton Cove State Park | TR | 201807 | Intertidal | 47.608 | -122.9856 |
| Twanoh State Park | TW | 201807 | Intertidal | 47.3785 | -122.9749 |
| P14 | NA | 201809 | Nearshore (WOAC) | 47.6067 | -122.9402 |
| P12 | NA | 201809 | Nearshore (WOAC) | 47.4253 | -123.108 |
| P11 | NA | 201809 | Nearshore (WOAC) | 47.3708 | -123.1333 |

Table S2: Complete list of potential HAB-forming ASV taxonomic assignments and COI sequences.

| Taxon | Sequence |
| --- | --- |
| Alexandrium_2b2 | ATTAAGCACTTCTTTCATGAGTTTATCACCTTCAAGTACAGGAAATCTTATCTTTGGATTATTAATCTCTGGTGTATCCTCATGTCTCACATCTCTTAACTTTTGGACAACAATTCTAAATCTGAGATCTTATTATCTGACATTAAAGACTATGCCATTATTCCCTTGGGCTCTCTTGATCACAGGAGGAATGCTTTTATTAACATTACCAATCTTATCAGGAGCTTTTCTAATGGTCTTGGCTGATCTTCATTCTAATACACTTTTCTTTGATCCAATCTTTGGAGGAGATCCTATATTCTATCAACACTTATTT |
| Alexandrium_3fc | ATTAAGCACTTCTTTCATGAGTTTATCACCTTCAAGTACAGGAAATCTTATCTTTGGATTATTAATCTCAGGTATATCCTCATGTCTCACATCTCTTAACTTTTGGACAACAATTCTAAATCTGAGATCTTATTATCTGACATTAAAGACTATGCCATTATTCCCTTGGGCTCTCTTGATTACAGGAGGAATGCTTTTATTAACATTACCAATCTTATCAGGAGCTTTTCTAATGGTCTTGGCTGATCTTCATTCTAATACACTTTTCTTTGATCCAATCTTTGGAGGAGATCCTATATTCTATCAACACTTATTT |
| Chaetoceros_00e | TTTATCTAGTGGTACCTCGCATTCAGGAGGTGCTGTTGATTTAGCTATTTTTAGTTTACACTTATCAGGAGCTTCATCAATTTTAGGTGCTATAAACTTTATTTGTACTATTTTTAATATGCGAGTTAAAAGTTTATCATTTCATAAGTTACCACTATTTGTATGGGCTGTATTAATAACAGCATTTTTACTTTTATTATCATTACCTGTTTTAGCGGGAGCTATTACAATGTTATTAACTGATAGAAATTTCAATACAACTTTTTTTGATCCAGCAGGTGGAGGTGATCCTATTTTATACCAACATTTATTT |
| Chaetoceros_163 | ATTATCTAGTGGAACCTCACACTCGGGTGGTGCTGTGGATTTAGCTATTTTTAGTTTACACTTATCTGGAGCTTCTTCTATTTTAGGAGCAATTAACTTTATTTGTACTATTTTTAATATGAGAGTAAAAAGTTTGGCTTTCCACAAGTTACCTTTGTTTGTTTGGGCAGTATTAATAACTGCGTTTTTATTATTACTATCGTTACCAGTATTAGCGGGAGCTATTACAATGTTATTAACTGACCGAAATTTTAACACCACATTTTTTGATCCTGCAGGGGGTGGTGACCCTGTATTATACCAGCACTTGTTT |
| Chaetoceros_17e | ATTATCTAGTGGAACCTCACACTCGGGTGGTGCTGTGGATTTAGCTATTTTTAGTTTACACTTATCTGGAGCTTCTTCTATTTTAGGAGCAATTAACTTTATTTGTACTATTTTTAATATGAGAGTTAAAAGTTTATCATTTCATAAATTACCTTTATTTGTATGGGCAGTGTTAATTACAGCATTTTTACTTTTATTATCACTACCAGTATTAGCAGGTGCTATTACAATGTTATTAACTGATAGAAATTTCAATACAACCTTTTTTGATCCAGCAGGAGGAGGTGACCCAATTTTATACCAACATTTATTT |
| Chaetoceros_211 | TTTATCAAGCGGTACTGCGCATTCAGGTGGAGCTGTTGATTTAGCAATTTTTAGTTTGCACTTATCAGGTGCTTCGTCAATTTTAGGAGCAATCAACTTTATTTGTACAATTTTTAATATGAGAGTTAAAAGTTTATCATTCCATAAATTACCTTTATTTGTTTGGTCTGTTTTAATAACAGCATTTTTACTTTTGTTATCTTTACCTGTTCTAGCTGGCGCTATTACTATGCTATTAACAGATAGAAATTTTAACACAACTTTTTTTGATCCAGCAGGTGGAGGTGATCCAGTTTTATACCAGCATTTATTT |
| Chaetoceros_4f5 | TTTATCAAGTGGTACATCACATTCAGGAGGTGCAGTAGATTTAGCTATTTTCAGTTTACACTTATCAGGAGCATCTTCTATTTTAGGTGCAATTAATTTTATATGTACTATTTTTAACATGCGTGTTAAAAGTTTATCGTTTCATAAACTTCCTTTATTTGTATGGGCTGTTTTAATTACAGCATTTTTATTATTGTTATCACTACCTGTTTTAGCTGGAGCAATTACAATGTTATTAACAGATCGAAATTTTAATACTACTTTCTTTGATCCAGCAGGTGGTGGAGACCCTGTATTATATCAACATTTATTC |
| Chaetoceros_559 | TCTATCTAGTATCACAGCCCACTCAGGTGGAGCCGTAGACTTAGCTATTTTTAGTTTACATGTTTCAGGAGCATCTTCAATTTTAGGTGCTATTAATTTTATTTGTACCATTTTTAACATGCGAGTAAAAAGTTTATCATTTCATAAACTTCCTTTATTTGTTTGGGCAGTTTTAATTACTGCATTTTTATTATTGTTATCTTTACCTGTTTTAGCTGGTGCGATTACTATGCTATTAACTGACCGGAATTTTAACACTACTTTTTTTGATCCTGCTGGTGGAGGTGACCCTGTTTTATATCAACATTTATTT |
| Chaetoceros_56d | TTTATCGAGCGGAACATCACATTCAGGCGGGGCTGTTGATTTAGCAATTTTTAGTTTACACTTATCAGGAGCATCTTCTATTTTAGGTGCAATAAACTTTATATGTACTATATTTAATATGCGTGTTAAAAGTTTATCGTTTCATAAATTACCTTTGTTTGTATGGGCTGTTTTAATAACAGCTTTTCTGTTATTACTATCATTACCTGTATTAGCAGGTGCGATTACAATGTTATTAACAGATAGAAATTTCAATACAACTTTTTTTGATCCAGCAGGTGGTGGAGATCCTGTTTTATATCAACATTTATTT |
| Chaetoceros_584 | TTTATCTAGCGGTACATCACACTCAGGTGGAGCTGTAGATTTAGCTATTTTTAGTTTACACTTATCTGGGGCTTCATCAATTTTAGGAGCTATTAACTTTATTTGTACTATTTTTAATATGAGAGTTAAAAGTTTATCATTTCATAAATTACCTTTATTTGTATGGGCAGTGTTAATTACAGCATTTTTACTTTTATTATCACTACCAGTATTAGCAGGTGCTATTACAATGTTATTAACTGATAGAAATTTCAATACAACCTTTTTTGATCCAGCAGGAGGAGGTGACCCAATTTTATACCAACATTTATTT |
| Chaetoceros_6c2 | TTTATCTAGTGGTACTGCACATTCAGGTGGAGCTGTAGATTTAGCTATTTTTAGCTTACATGTTTCTGGAGCATCTTCAATATTAGGAGCTATTAATTTTATTTGTACTATTTTCAATATGAGAGTAAAAAGTTTATCATTCCATAAACTTCCTTTATTTGCTTGGTCAGTTTTAATCACTGCTTTTTTATTATTATTATCTTTACCAGTTTTAGCAGGAGCTATCACTATGCTTTTAACAGATAGAAATTTTAATACTACTTTTTTTGATCCCGCAGGTGGAGGTGATCCAATTTTATACCAACATTTATTT |
| Chaetoceros_734 | TTTATCAAGTGGTACATCACATTCAGGAGGTGCAGTAGATTTAGCTATTTTCAGTTTACACTTATCAGGAGCATCTTCTATTTTAGGTGCAATTAATTTTATATGTACTATTTTTAACATGCGTGTTAAAAGTTTATCGTTTCATAAACTTCCTTTATTTGTATGGGCTGTTTTAATTACAGCATTTTTATTATTGTTATCACTACCTGTTTTAGCTGGAGCAATTACGATGTTATTAACAGATCGAAATTTTAATACTACTTTCTTTGATCCAGCAGGTGGTGGAGACCCTGTATTATATCAACATTTATTC |
| Chaetoceros_788 | TTTATCAAGCGGTACTTCGCATTCAGGTGGTGCCGTTGACTTGGCTATTTTTAGTCTACACCTTTCAGGAGCTTCTTCGATTTTAGGTGCTATTAATTTTATTTGTACAATTTTTAACATGAGAGTAAAAAGTCTTTCTTTTCATAAATTACCTTTATTTGTATGGGCAGTTTTAATTACAGCGTTTTTACTTCTTTTATCATTGCCTGTTCTAGCGGGCGCTATCACAATGCTATTAACTGATAGAAATTTCAATACAACCTTTTTTGATCCAGCAGGAGGAGGTGACCCAATTTTATACCAACATTTATTT |
| Chaetoceros_83a | TTTATCTAGCGGAACTTCGCATTCAGGTGGGGCTGTTGATTTAGCAATTTTTAGTTTACACTTATCAGGAGCATCTTCTATTTTAGGTGCAATAAACTTTATATGTACTATATTTAATATGCGTGTTAAAAGTTTATCGTTTCATAAATTACCTTTGTTTGTATGGGCCGTTTTAATAACAGCTTTTCTGTTATTACTATCATTACCCGTATTAGCAGGTGCAATTACGATGCTATTAACAGATAGAAATTTTAATACAACTTTTTTTGATCCTGCAGGTGGTGGAGATCCTGTTCTATATCAGCATTTATTT |
| Chaetoceros_a8e | TCTATCTAGTATCACAGCCCACTCAGGTGGAGCCGTAGACTTAGCTATTTTTAGTTTACATGTTTCAGGAGCATCTTCAATTTTAGGTGCTATTAATTTTATTTGTACCATTTTTAACATGCGAGTAAAAAGTTTATCATTTCATAAACTTCCTTTATTTGTTTGGGCAGTTTTAATTACTGCATTTTTATTATTGTTATCTTTACCTGTTTTAGCTGGTGCGATTACTATGCTATTAACTGACCGTAATTTTAACACTACTTTTTTTGATCCTGCTGGTGGAGGTGACCCTGTTTTATATCAACATTTATTT |
| Chaetoceros_ab3 | TTTATCTAGTGGTACTTCACATTCAGGAGGTGCTGTTGATTTAGCTATTTTTAGTTTACACTTATCAGGAGCTTCATCAATTTTAGGTGCTATAAACTTTATTTGTACAATCTTTAACATGCGAGTTAAAAGTTTATCGTTCCATAAATTACCACTATTTGTATGGGCGGTATTAATAACAGCATTTTTACTTTTATTGTCATTACCTGTTTTAGCAGGCGCTATTACAATGTTATTAACAGATAGAAATTTCAATACAACATTCTTTGATCCTGCTGGTGGAGGTGACCCAATTTTATACCAACATTTATTT |
| Chaetoceros_ae1 | TTTATCCAGTGGAACTGCTCATTCAGGAGGTGCTGTTGATTTAGCTATTTTTAGTTTACATTTATCAGGAGCATCTTCTATTTTGGGAGCTATAAATTTTATATGCACAATTTTTAATATGCGAGTTAAAAGTTTATCATTTCATAAATTGCCTTTATTCGTTTGGTCAGTTTTGATCACAGCTTTCTTGCTTCTTTTATCACTACCTGTTTTAGCCGGTGCTATTACTATGTTATTAACAGATCGTAATTTTAACACTACCTTTTTTGACCCTGCCGGTGGAGGTGATCCTGTTTTATATCAACATTTATTT |
| Chaetoceros_b14 | TTTATCTAGTGGTACTGCACATTCAGGTGGAGCTGTAGATTTAGCTATTTTTAGCTTACATGTTTCTGGAGCATCTTCAATATTAGGAGCTATTAATTTTATTTGTACTATTTTCAATATGAGAGTAAAAAGTTTATCATTCCATAAACTTCCTTTATTTGCTTGGTCAGTTTTAATCACTGCTTTTTTATTATTATTATCTTTACCAGTTTTAGCAGGAGCTATCACTATGCTTTTAACAGATAGAAATTTTAATACTACTTTTTTTGATCCTGCAGGTGGAGGTGATCCAATTTTATACCAACATTTATTT |
| Chaetoceros_b25 | TTTATCAAGCGGAACTTCTCATTCAGGAGGTGCTGTTGATTTAGCTATATTTAGTTTACATCTATCAGGAGCATCATCAATTTTAGGAGCTATAAATTTTATTTGTACAATTTTTAACATGAGAGTTAAAAGTTTATCTTTTCATAAATTACCGTTATTTGTTTGGTCAGTTTTAATTACAGCATTTTTATTATTACTTTCTTTACCTGTGTTGGCGGGAGCAATAACAATGCTATTAACTGATAGAAATTTTAACACTACCTTTTTTGACCCTGCGGGTGGAGGTGACCCTATATTATATCAACATTTATTT |
| Chaetoceros_c4f | TTTATCTAGTGGTACCTCGCATTCAGGAGGTGCTGTTGACTTAGCTATTTTTAGTTTACATTTATCAGGAGCTTCATCAATTTTAGGTGCTATAAACTTTATTTGTACTATTTTTAATATGCGAGTTAAAAGTTTATCATTTCATAAGTTACCACTATTTGTATGGGCTGTATTAATAACAGCATTTTTACTTTTATTATCATTACCTGTTTTAGCGGGAGCTATTACAATGTTATTAACTGATAGAAATTTCAATACAACTTTTTTTGATCCAGCAGGTGGAGGTGATCCTATTTTATACCAACATTTATTT |
| Chaetoceros_cc3 | TTTATCTAGTGGTACTGCACATTCAGGTGGAGCTGTAGATTTAGCTATTTTTAGCTTACATGTTTCTGGAGCATCTTCAATATTAGGAGCTATTAATTTTATTTGTACTATTTTCAATATGAGAGTAAAAAGTTTATCATTCCATAAACTTCCTTTATTTGCTTGGTCAGTTCTAATCACTGCATTTTTATTATTATTATCTTTACCAGTTTTAGCAGGAGCTATCACTATGCTTTTAACAGATAGAAATTTTAATACTACTTTTTTTGATCCCGCAGGTGGAGGTGATCCAATTTTATACCAACATTTATTT |
| Chaetoceros_dcd | TTTATCAAGTGGAACTTCACATTCAGGTGGAGCTGTTGATTTAGCAATTTTTAGTTTACACTTATCAGGAGCATCTTCTATTTTAGGTGCAATAAACTTTATATGTACTATATTTAATATGCGTGTTAAAAGTTTATCGTTTCATAAATTACCTTTGTTTGTATGGGCTGTTTTAATAACAGCTTTTCTGTTATTACTATCATTACCTGTATTAGCAGGTGCGATTACAATGTTATTAACAGATAGAAATTTTAATACGACTTTTTTTGATCCTGCAGGTGGTGGAGATCCTGTTTTATATCAACATTTATTT |
| Chaetoceros_e58 | TTTATCTAGTGGTACTGCACATTCAGGTGGAGCTGTAGATTTAGCTATTTTTAGCTTACATGTTTCTGGAGCATCTTCAATATTAGGAGCTATTAATTTTATTTGTACTATTTTCAATATGAGAGTAAAAAGTTTATCATTCCATAAACTTCCTTTATTTGCTTGGTCAGTTTTAATCACTGCATTCTTATTATTATTATCTTTACCAGTTTTAGCAGGAGCTATCACTATGCTTTTAACAGATAGAAATTTTAATACTACTTTTTTTGATCCCGCAGGTGGAGGTGATCCAATTTTATACCAACATTTATTT |
| Chaetoceros_ed1 | ATTATCTAGTGGTACTTCTCACTCAGGAGGGGCTGTTGATTTAGCAATTTTTAGTTTACACTTATCAGGAGCGTCTTCTATTTTAGGCGCTATTAATTTTATTTGTACAATTTTTAATATGCGGGTAAAGAGTCTTGCATTTCACAAATTACCTTTATTTGTGTGGGCTGTTTTAATCACAGCGTTTCTATTATTACTTTCTTTACCAGTTTTAGCAGGAGCAATTACAATGCTGTTAACTGATAGAAATTTCAATACAACCTTTTTTGACCCTGCAGGAGGAGGAGATCCTGTTTTATACCAGCACTTATTT |
| Chaetoceros_edb | ATTATCAAGTGGTACAGCACATTCAGGTGGAGCTGTAGATTTAGCTATTTTTAGTTTACATGTTTCTGGAGCATCTTCAATATTAGGAGCTATTAATTTTATTTGTACTATCTTTAATATGAGAGTAAAAAGTTTATCATTTCATAAACTTCCGTTATTTGCTTGGTCAGTTTTAATTACTGCATTTTTATTATTGTTATCTTTACCAGTTTTAGCGGGAGCTATTACTATGCTTTTAACTGATAGAAATTTTAATACTACTTTCTTTGATCCTGCAGGTGGAGGTGATCCAATTTTATACCAACATTTATTT |
| Chaetoceros_efa | TTTATCGAGTGGTACATCACATTCAGGTAGTGCCGTAGATTTAGCTATTTTTAGTTTACATATTTCAGGAGCTTCTTCTATTTTAGGTGCAATTAATTTTATTTGTACCATTTTTAATATGCGAGTTAAAAGTTTATCATTCCATAAATTACCATTATTTGTATGGTCTGTTTTAATAACAGCATTTTTACTACTATTATCTTTACCAGTTTTAGCAGGTGCAATCACTATGTTATTAACTGATAGAAATTTCAATACTACATTTTTTGATCCTGCTGGAGGAGGTGACCCTGTTTTATATCAACATTTATTT |
| Chattonella_135 | ATTAAGTAGTGTTCAAGCACACTCTGGGCCTTCAGTTGACTTAGCAATCTTTAGTCTTCACTTATCGGGGGCTGCATCAATTTTAGGAGCAATAAACTTTATTACTACTATTTTTAATATGCGAGCACCAGGTATGACAATGCATAGATTACCACTGTATGTTTGGTCTATTTTAATTACTTCATTCCTTTTACTTCTTTCTTTACCCGTATTAGGAGGAGCAATTACTATGTTATTAACTGATAGAAATTTTAATACTTCATTCTTTGATCCAGCTGGTGGAGGTGATCCAATTTTATTTCAACATTTATTT |
| Chattonella_13c | ATTAAGTAGTGTTCAAGCACACTCTGGGCCTTCAGTTGACTTAGCAATCTTTAGTCTTCACTTATCGGGGGCTGCATCAATTTTAGGGGCAATAAACTTTATTACTACCATCTTTAATATGCGCGCACCGGGTATGACAATGCATAGATTACCACTGTATGTTTGGTCTATTTTAATTACTTCATTCCTTTTACTTCTGTCTTTACCCGTATTAGGAGGAGCAATTACTATGTTATTGACTGATAGAAATTTTAATACTTCATTCTTTGACCCAGCTGGTGGAGGAGATCCAATTTTATTTCAACATTTATTT |
| Chattonella_e0f | ATTAAGTAGTGTTCAAGCACACTCTGGGCCTTCAGTTGACTTAGCAATCTTTAGTCTTCACTTATCGGGGGCTGCATCAATTTTAGGGGCAATAAACTTTATTACTACCATCTTTAATATGCGCGCACCGGGTATGACAATGCATAGGTTACCACTGTATGTTTGGTCTATTTTAATTACTTCATTCCTTTTACTTCTGTCTTTACCCGTATTAGGAGGAGCAATTACTATGTTATTGACTGATAGAAATTTTAATACTTCATTCTTTGACCCAGCTGGTGGAGGAGATCCAATTTTATTTCAACATTTATTT |
| Chrysochromulina_18f | ATTAGCTAGTATTCAAGCACATTCAGGTGGTTCTGTTGATTGTGCTATTTTTTCATTACATATTGCTGGTGTATCTTCTATTTTAGGTGCTATTAATTTTATTGTTACAATTAGTAATATGCGTGCACCAGGAATGACTGCAAATAGAACACCTTTATTTGTTTGAGCTGTTTTTATTACTGCTTTTTTACTTTTATTATCATTACCGGTTTTAGCTGGTGCAATAACAATGTTATTGACTGATCGTAATTTTAATACATCTTTTTTTGATCCAAACGGTGGTGGTGATCCTGTTTTATACCAACATTTATTT |
| Chrysochromulina_34d | ATTGTCAGGAATTCAAGCACATTCCGGTGGTTCTGTTGATTGTGCTATTTATTCGCTTCATCTAGCTGGTGTTTCTTCAATTTTAGGAGCAATCAACTTCATTGTTACAATCACTAACATGCGTGCTCCAGGAATGACTGCTAATCGAACTCCTTTGTTCGTTTGAGCTGTTTATATTACAGCATTTTTACTTTTGCTTTCTTTACCGGTGCTTGCTGGTGCAATCACAATGTTACTGACAGACCGTAATTTTAACACCTCTTTTTTTGATCCAAACGGTGGAGGTGATCCCGTTTTGTACCAACACCTGTTC |
| Chrysochromulina_75f | TTTAGCTGGTATTCAAGCTCATTCAGGTGGTTCTGTTGACTGTGCTATTTATTCACTTCACTTAGCAGGTGTTTCTTCAATTTTAGGTGCTATTAATTTTATTGTTACAATTACAAATATGCGTGCACCTGGAATGACTGCAAACCGTACACCTCTTTTTGTTTGAGCAGTATACATTACTGCATTCTTACTATTACTTTCTTTACCTGTACTTGCAGGAGCAATTACAATGTTACTTACAGATCGTAACTTTAATACTTCTTTCTTTGATCCTAACGGTGGTGGTGATCCTGTTTTATACCAACACTTGTTT |
| Chrysochromulina_7aa | TTTGTCTGGCATTCAAGCTCATTCAGGCGGTTCTGTTGATTGCGCAATTTACTCGTTGCATTTAGCCGGTGTTTCTTCAATTTTAGGCGCAATTAATTTTATTGTAACAATAACTAATATGCGTGCACCTGGACTATCTGCTAATCGAACGCCTCTTTTTGTTTGAGCTGTTTATATTACAGCGTTTTTATTGTTACTTTCTTTACCCGTTTTAGCTGGTGCAATTACGATGTTGCTAACTGATCGTAATTTTAACACTTCATTCTTTGATGCGAATGGCGGTGGTGACCCTGTATTGTATCAACACTTATTT |
| Dinophysis_2bc | ATTAAGCACTTCTTTCTTGAGTTTATCACCTTCAAGTACAGGAAATCTTATCTTTGGATTATTAATCTCAGGAATCTCCTCATGTCTCACATCTCTTAACTTTTGGACAACAATTTGAAATCTGAGATCTTATTACTTAACATTAAAGACTATGCCATTATTCCTTTGGTCTCTCTTGATTACAGGAGGAATGCTTTTATTAACATTACCAATCTTATCAGGAGCTCTTCTAATGGTCACAGCTGATCTTCATTCTAATACACTTTTCTTTGATCCAATCTTTGAAGGAGATCCTATATTCTATCAACACTTATTT |
| Dinophysis_2d8 | ATTAAGCACTTCTTTCTTGAGTTTATCACCTTCAAGTACAGGAAATCTTATCTTTGGATTATTAATCTCAGGAATCTCCTCATGTCTCACATCTCTTAACTTTTGGACAACAATTATAAATCTGAGATCTTATTACTTAACATTAAAGACTATGCCATTATTCCTTTGGTCTCTCTTGATTACAGGAGGAATGCTTTTATTAACATTACCAATCTTATCAGGAGCTCTTCTAATGGTCACAGCTGATCTTCATTCTAATACACTTTTCTTTGATCCAATCTTTGAAGGAGATCCTATATTCTATCAACACTTATTT |
| Dinophysis_a88 | ATTAAGCACTTCTTTCTTGAGTTTATCACCTTCAAGTACAGGAAATCTTATCTTTGGATTATTAATCTCAGGAATCTCCTCATGTCTCACATCTCTTAACTTTTGGACAACAATTCTAAATCTGAGATCTTATTACTTAACATTAAAGACTATGCCATTATTCCTTTGGTCTCTCTTGATTACAGGAGGAATGCTTTTATTAACATTACCAATCTTATCAGGAGCTCTTCTAATGGTCACAGCTGATCTTCATTCTAATACACTTTTCTTTGATCCAATCTTTGAAGGAGATCCTATATTCTATCAACACTTATTT |
| Gonyaulax_564 | ATTAAGCACTTCTTTCATGAGTTTATCACCTTCAAGTTTGGGAAATCTTATCTTTGGATTATTAATCTCAGGTATATCCTCATGTCTCACATCTCTTAACTTTTGGACAACAATTCTAAATCTGAGATCTTATTATCTGACATTAAAGACTATGCCATTATTCCCTTGAGCTCTCTTGATTACAGGAGGAATGCTTTTATTAACATTACCAATCTTATCAGGAGCTCTTCTAATGGTCTTGGCTGATCTTCATTCTAATACACTTTTCTTTGATTCAATCTTTGGAGGAGATCCTATATTCTATCAACACTTATTT |
| Gymnodinium_e25 | ATTAAGCACTTCTTTCATGGGTTTATCACCTTCAAGTACAGCTTTCATGGTCTTTGGATTATTAATGTCAGGTATATCCTCATCTCTCACATCTCTTAACTTTTGGACAACAATTCTAAATCTGAGATCTTATTATCTGTCATTAAAGACTATACCATTATTCCCTTGGGCTCTCTTGATAACAGGAGGAATGCTTTTATTAACATTACCAATCTTATCTGGAGCTCTTCTAATGGTCTTGGCTGATATTCATTCTAATACACTTTTCTTTGATCCAATCTTTGGAGGTGATCCTATATTCTATCAACACTTATTT |
| Gymnodinium_eb3 | ATTAAGCACTTCTTTCATGAGTTTATCACCTTCAAGTACAGCTTTCATGGTCTTTGGATTATTAATGTCAGGTATATCCTCATCTCTCACATCTGTTAACTTTTGGACAACAATTCTAAATCTGAGATCTTATTATCTGTCATTAAAGACTATACCATTATTCCCTTGGGCTCTCTTGATAACAGGAGGAATGCTTTTATTAACATTACCAATCTTATCTGGAGCTCTTCTAATGGTCTTGGCTGATCTTCATTCTAATACACTTTTCTTTGATCCAATCTTTGGAGGTGATCCTATATTCTATCAACACTTATTT |
| Hematodinium_19f | ATTAAGTACATCATTAATAAGTCTATCACCTATTGGAATTGATATTTTATTATATGGATTATTATTGTCAGGTATATCATCATGTCTAACATCTATTAATTTTATCGCTACAATTATAAATATGAGATGTTATAGTATGACATTATCGATTATGCCAGTATATACATGGTCTATAAATATTACAGGATTTCTATTGTTATTAACATTACCTATATTAACAGGAGCTCTTATAATGTCGTTAGCAGATCTTCATTATAATACAGTTTTCTTTAATCCAATATTTGGAGGCGATCCTGTACTTTATCAACATTTATTT |
| Hematodinium_43a | ATTAAGTACATCATTAATAAGTCTATCACCTATTGGAATTGATATTTTATTATATGGATTATTATTGTCAGGTATATCATCATGTCTAACATCTATTAATTTCATCGCTACAATTATAAATATGAGATGTTATAGTATGACATTATCGATTATGCCAGTATATACATGGTCTATAAATATTACAGGATTTCTATTGTTATTAACATTACCTATATTAACAGGAGCTCTTATAATGTCGTTAGCAGATCTTCATTATAATACAGTTTTCTTTAATCCAATATTTGGAGGCGATCCTGTACTTTATCAACATTTATTT |
| Hematodinium_449 | ATTAAGTACATCATTAATAAGTCTATCACCTATTGGAATTGATATTTTATTATATGGATTATTATTGTCAGGTATATCATCATGTCTAACATCTATTAATTTCATCGCTACAATTATAAATATGAGATGTTATAGTATGACATTATCGATTATGCCAGTATATACATGGTCTATAAATATTACAGGATTTCTATTGTTATTAACATTACCTATATTAACAGGAGCTCTTATAATGTCGTTAGCAGATCTTCATTATAATACAGTTTTCTTTAATCCAATATTTGGAGGCGATCCTGTACTTTATCAACATTTCTTT |
| Hematodinium_a88 | ATTAAGTACATCATTAATAAGTCTATCACCTATTGGAATTGATATTTTATTATATGGATTATTATTGTCAGGTATATCATCATGTCTAACATCTATTAATTTCATCGCTATAATTATAAATATGAGATGTTATAGTATGACATTATCGATTATGCCAGTATATACATGGTCTATAAATATTACAGGATTTCTATTGTTATTAACATTACCTATATTAACAGGAGCTCTTATAATGTCGTTAGCAGATCTTCATTATAATACAGTTTTCTTTAATCCAATATTTGGAGGCGATCCTGTACTTTATCAACATTTATTT |
| Heterocapsa_2f5 | TCTATCTACTTCTTTCCTATCTCTATCCCCATCTTCTATGTACTTCCTACTATCTGGTCTACTAGTTTCTGGTCTATCTTCTGCTCTAACTTCTCTAAACTTCTTCCTAACAATTCTTAACATGCGTTGTTTCTCTATGAATCTAAAGCTACTACCTCTATTCAACTGGTCTATCCTAATTACTTCTGTCCTACTTCTATTCACTCTACCTGTTCTATCTGGTGCTGTCGTTATGATCCTATCCGATCTATCTGCTAATACACTATTCTATGATCCAATCTTCGGTGGTGATCCTGTTCTTTACCAGCATCTTTTC |
| Heterocapsa_49f | TCTATCTACTTCTTTCCTATCTCTATCTCCATCTTCTATGTATTTCCTACTATCTGGTCTACTAGTTTCTGGTCTATCTTCTGCTCTAACTTCTCTAAACTTCTTCCTAACAATTCTTAACATGCGTTGTTTCTCTATGAATCTAAAGCTACTACCTCTATTCAACTGGTCTATCCTAATTACTTCTGTCCTACTTCTACTAACTCTACCTGTTCTATCTGGTGCTGTCGTTATGATCCTATCCGATCTATCTGCTAATACACTATTCTATGATCCAATCTTCGGTGGTGATCCTGTTCTTTACCAGCATCTTTTC |
| Heterocapsa_56d | TCTATCTACTTCTTTCCTATCTCTATCCCCATCTTCTATGTATTTCCTACTATCTGGTCTACTAGTTTCTGGTCTATCTTCTGCTCTAACTTCTCTAAACTTCTTCCTAACAATTCTTAACATGCGTTGTTTCTCTATGAATCTAAAGCTACTACCTCTATTCAACTGGTCTATCCTAATTACTTCTGTCCTACTTCTACTAACTCTACCTGTTCTATCTGGTGCTGTCGTTATGATCCTATCCGATCTATCTGCTAATACACTATTCTATGATCCAATCTTCGGTGGTGATCCTGTTCTTTACCAACATCTTTTC |
| Heterocapsa_7b2 | TCTATCTACTTCTTTCCTATCTCTATCCCCATCTTCTATGTACTTCCTACTATCTGGTCTACTAGTTTCTGGTCTATCTTCTGCTCTAACTTCTCTAAACTTCTTCCTAACAATTCTTAACATGCGTTGTTTCTCTATGAATCTAAAGCTACTACCTCTATTCAACTGGTCTGTTCTAATTACTTCTGTCCTACTTCTATTCACTCTACCTGTTCTATCTGGTGCTGTCGTTATGATCCTATCCGATCTATCTGCTAATACACTATTCTATGATCCAATCTTCGGTGGTGATCCTGTTCTTTACCAGCATCTTTTC |
| Heterocapsa_995 | TCTATCTACTTCTTTCCTATCTCTATCCCCATCTTCTATGTATTTCCTACTATCTGGTCTACTAGTTTCTGGTCTATCTTCCGCATTAACATCTCTAAATTTCTTCCTAACAATCCTTAACATGCGTTGCTTCTCTATGAATCTAAAACTTCTACCTCTATTTAACTGGTCTATTATTATTACTTCTGTTCTACTACTATTCACTCTACCTGTCCTATCTGGTGCAGTTGTTATGATCCTTTCTGATCTTTCTTGCAATACTCTATTCTATGACCCTATTTTCGGTGGTGATCCAGTCCTATACCAACACCTTTTC |
| Heterocapsa_adc | TCTATCTACTTCTTTCCTATCTCTATCCCCATCTTCTATGTATTTCCTACTATCTGGTCTACTAGTTTCTGGTCTATCTTCTGCTCTAACTTCTCTAAACTTCTTCCTAACAATTCTTAACATGCGTTGTTTCTCTATGAATCTAAAGCTACTACCTCTATTCAACTGGTCTATCCTAATTACTTCTGTCCTACTTCTACTAACTCTACCTGTTCTATCTGGTGCTGTCGTTATGATCCTATCCGATCTATCTGCTAATACACTATTCTATGATCCAATCTTCGGTGGTGATCCTGTTCTTTACCAGCATCTTTTC |
| Heterocapsa_ba5 | TCTATCTACTTCTTTCCTATCTCTATCTCCATCTTCTATGTATTTCCTACTATCTGGTCTACTAGTTTCTGGTCTATCTTCTGCTCTAACTTCTCTAAACTTCTTCCTAACAATTCTTAACATGCGTTGTTTCTCTATGAATCTAAAGCTACTACCTCTATTCAACTGGTCTATCCTAATTACTTCTGTCCTACTTCTACTAACTCTACCTGTTCTATCTGGTGCTGTCGTTATGATCCTATCCGATCTATCTGCTAATACACTATTCTATGATCCAATCTTCGGTGGTGATCCTGTTCTTTACCAACATCTTTTC |
| Heterocapsa_ddc | TCTATCTACTTCTTTCCTATCTCTATCCCCATCTTCTATGTATTTCCTACTATCTGGTCTACTAGTTTCTGGTCTATCTTCCGCATTAACATCTCTAAATTTCTTCCTAACAATCCTTAACATGCGTTGCTTCTCTATGAATCTAAAACTTCTACCTCTATTTAACTGGTCTATTATTATTACTTCTGTTCTACTACTATTCACTCTACCTGTCCTATCTGGTGCAGTCGTTATGATCCTTTCTGATCTTTCTTGCAATACTCTATTCTATGACCCTATTTTCGGTGGTGATCCAGTCCTATACCAACACCTTTTC |
| Heterosigma_c9e | ATTAAGTAGCGCTCAAGCTCACTCAGGACCGTCGGTAGATTTAGCTATTTTCAGTTTACACGTTTCAGGAGCAGCATCAATTTTAGGGGCAATTAATTTTATTACCACTATTTTAAACATGCGAGCACCTGGTATGACCATGCATCGACTACCGTTGTTTGTGTGGGCTGTGTTTATTACTGCAATTTTATTATTATTATCGTTACCAGTATTAGCAGGAGCAATTACTATGTTATTAACTGATCGAAATTTCAACACTACCTTTTACGATCCGGCAGGAGGAGGAGACCCTGTATTGTATCAACATTTATTT |
| Karlodinium_057 | ATTAAGTACTTCTTTCATGAGTTTATCACCTTCAACTACAGCTTATCTTATCTTTGGATTATTAATGTCAGGTATATCTTCATGTCTAACTTCTATTAACTTTTTTATTACAATTCTTAATCTGAGATCTTATTATCTTACATTAAAGACTATGCCATTATTCCCATGGTCTCTTTTGATAACAGGAGGAATGCTATTATTAACATTACCAATCTTATCAGGTGCATTACTAATGGTCTTGGCTGATCTTCATTGTAATTCATCATTCTTTGATCCAATCTTTGGAGGAGATCCTATATTCTATCAACATTTATTT |
| Karlodinium_69b | ATTAAGCACTTCTTTCATGAGTTTATCACCTTCAACCACAGCTTATCTTATCTTTGGATTATTAATGTCAGGTATATCCTCATGTCTCACTTCTATTAACTTTTTTATTACAATTCTTAATCTGAGATCTTATTATCTTACATTAAAGACTATGCCATTATTCCCATGGTCTCTCTTGATAACAGGAGGAATGCTATTATTAACATTACCAATCTTATCAGGTGCATTCCTAATGGTCTTGGCTGATCTTCATTCTAATTCACTTTTCTTTGATCCAATCTTTGGAGGAGATCCTATATTCTATCAACACTTATTT |
| Karlodinium_8ed | ATTAAGCACTTCTTTCATGAGTTTATCACCTTCAACTACAGCTTATCTTATCTTTGGATTATTAATGTCAGGTATATCCTCATGTCTAACTTCTATTAACTTTTTTATTACAATTCTTAATCTGAGATCTTATTATCTTACATTAAAGACTATGCCATTATTCCCATGGTCTCTTTTGATAACAGGAAGAATGCTATTATTAACATTACCAATCTTATCAGGTTCTCTATTAATGGTCTCTGCTGATCTTCATTGTAATTCATCATTCTTTGATCCAATCTTTGGAGGAGATCCTATATTCTATCAACATTTATTT |
| Karlodinium_a27 | ATTAAGCACTTCTTTCATGAGTTTATCACCTTCAACTACAGCTTATCTTATCTTTGGATTATTAATGTCAGGTATATCCTCATGTCTAACTTCTATTAACTTTTTTATTACAATTCTTAATCTGAGATCTTATTATCTTACATTAAAGACTATGCCATTATTCCCATGGTCTCTTTTGATAACAGGAGGAATGCTATTATTAACATTACCAATCTTATCAGGTGCATTACTAATGGTCTTGGCTGATCTTCATTGTAATTCACTTTTCTTTGATTCAATCTTTGGAGGAGATCCTATATTCTATCAACATTTATTT |
| Karlodinium_abe | ATTAAGCACTTCTTTCATGAGTTTATCACCTTCAACTACAGCTTATCTTATCTTTGGATTATTAATGTCAGGTATATCCTCATGTCTAACTTCTATTAACTTTTTTATTACAATTCTTAATCTGAGATCTTATTATCTTACATTAAAGACTATGCCATTATTCCCATGGTCTCTTTTGATAACAGGAGGAATGCTATTATTAACATTACCAATGTTATCAGGTGCATTACTAATGGTCTTGGCTGATCTTCATTCTAATTCATCATTCTTTGATCCAATCTTTGGAGGAGATCCTATATTCTATCAACATTTATTT |
| Nitzschia_010 | TTTATCAGGAATTATCGCTCACTCAGGAGGTGCTGTTGATTTAGCTATTTTCAGTTTACACCTTTCAGGTGCTGCGTCTATTCTAGGTGCTATTAATTTCATCTGTACTATTGTGAACATGAGAACTGAAAGCTTACCATTTCACAAATTACCTTTATTTGTTTGGTCAGTGTTTTTAACAGCAATTCTTTTATTATTATCTCTACCAGTGTTAGCAGGTGCTATTACAATGTTATTAACTGATAGAAATTTCAATACAACATTCTTTGATCCAGCAGGTGGTGGTGATCCAGTTCTTTACCAGCACTTATTT |
| Nitzschia_065 | TTTATCAGGTATTATTGCACACTCTGGTGGTGCCGTTGATTTAGCTATTTTCAGTTTACACCTTTCAGGTGCTGCTTCTATTTTAGGTGCGATTAATTTTATTTGTACAATTGTAAACATGAGAACCGAGAGTTTACCATTCCACAAATTACCTTTATTTGTATGGTCAGTGTTTTTAACAGCAATTCTTCTATTATTATCTTTACCTGTACTAGCAGGTGCTATTACAATGCTACTGACTGATAGAAATTTTAATACAACTTTTTTTGATCCAGCAGGTGGTGGTGATCCAGTTCTTTACCAGCATTTGTTT |
| Nitzschia_089 | CCTTTCAGGTATAATAGCTCATTCAGGTGGTGCTGTTGATTTGGCTATTTTCAGTTTACACCTTTCAGGTGCTGCATCTATTCTAGGTGCAATTAATTTCATCTGTACTATTGTAAACATGAGAACTGAAAGTTTACCATTTCATAAGTTACCATTATTTGTATGGTCAGTTTTTTTAACAGCAATTTTATTATTACTGTCTTTACCAGTATTAGCAGGTGCTATTACTATGTTATTGACTGATAGAAATTTCAACACTACTTTTTTCGACCCAGCAGGTGGTGGTGATCCAGTTCTTTACCAGCATTTGTTT |
| Nitzschia_0c1 | TTTATCTGGTATTATTGCTCACTCTGGTGGTGCTGTTGATCTTGCAATTTTTAGTTTACATTTATCAGGAGCTGCTTCGATTTTAGGAGCAATTAATTTTATTTGTACTATTGTAAATATGCGAACTGACAGTTTACCTTTCCATAAATTACCGTTGTTTGTGTGGGCAGTTTTCATTACAGCTATTCTTCTACTATTGTCTCTACCTGTTTTAGCTGGTGCCATAACTATGTTACTGACAGATCGGAATTTTAATACTACTTTTTTTGACCCTGCAGGTGGTGGTGATCCAGTTTTATATCAGCATTTATTC |
| Nitzschia_2bb | TTTATCAGGTATTATTGCACACTCTGGTGGTGCCGTTGATTTAGCTATTTTCAGTTTACACCTTTCAGGTGCTGCTTCTATTTTAGGTGCGATTAATTTTATTTGTACAATTGTAAACATGAGAACCGAGAGTTTACCATTCCACAAATTACCTTTATTTGTATGGTCAGTGTTTTTAACAGCAATTCTTCTATTATTATCTTTGCCTGTACTAGCAGGTGCTATTACAATGTTACTGACTGATAGGAATTTTAATACAACTTTTTTTGATCCAGCAGGTGGTGGTGATCCAGTTCTTTACCAGCATTTGTTT |
| Nitzschia_2da | TTTATCAGGTATTATTGCACACTCTGGTGGTGCTGTTGATTTAGCTATTTTCAGTTTACACCTTTCAGGTGCTGCTTCTATTTTAGGTGCGATTAATTTTATTTGTACAATTGTAAACATGAGAACTGAGAGTTTACCATTCCACAAATTACCTTTATTTGTGTGGTCGGTGTTTTTAACAGCAATCCTTCTATTATTATCTTTACCTGTACTAGCAGGTGCTATTACAATGTTACTGACTGATAGAAATTTTAATACAACTTTTTTTGATCCAGCAGGTGGTGGTGATCCAGTTCTTTACCAGCATTTGTTT |
| Nitzschia_371 | TTTATCAGGTATTATTGCACACTCTGGTGGTGCTGTTGATTTAGCTATTTTTAGTTTACACCTTTCAGGTGCTGCTTCTATTTTAGGTGCGATTAATTTTATTTGTACAATTGTAAACATGAGAACTGAGAGTTTACCATTCCACAAATTACCTTTATTTGTATGGTCAGTGTTTTTAACAGCAATTCTTCTATTATTATCTTTACCTGTACTAGCAGGTGCTATTACGATGTTACTGACTGATAGAAATTTTAATACAACTTTTTTTGATCCAGCAGGTGGTGGTGATCCAGTTCTTTACCAGCATTTGTTT |
| Nitzschia_456 | ACTGTCGGGTGTTATCGCACACTCGGGAGGTTCGGTAGACTTAGCAATTTTCAGTCTTCACTTATCTGGAGCTGCGTCTATCTTAGGTGCAATTAATTTCATTTGTACTATTGTAAACATGCGAACAGAAAGCTTACCTTTCCATAAGCTACCTTTGTTTGTATGGTCTGTTTTCATTACTGCTATTTTATTATTATTATCGTTACCGGTATTAGCAGGAGCTATTACAATGCTGCTTACAGATCGAAATTTCAACACTACTTTTTTTGATCCAGCAGGTGGTGGAGATCCTGTATTATTCCAGCACTTATTC |
| Nitzschia_5b5 | TTTATCAAGTATAACAGCACATTCAGGAGGATCTGTAGATTTAGCTATTTTCAGTCTTCATTTAGCAGGAGCTTCTTCTATTTTAGGAGCCATTAATTTTATTTGTACTATTGTTAACATGCGTACAGACAGTTTACCATTTCATAAATTACCTTTATTTGTTTGGTCTGTTCTTATTACTGCTGTATTATTACTATTATCTTTACCTGTTTTAGCAGGTGCAATTACAATGTTACTAACTGATAGAAATTTTAATACAACTTTTTTTGATCCAGCAGGTGGAGGTGATCCAGTTCTTTATCAGCATTTGTTT |
| Nitzschia_616 | TTTATCAGGTATTATTGCACACTCTGGTGGTGCCGTTGATTTAGCTATTTTCAGTTTACACCTTTCAGGTGCTGCTTCTATTTTAGGTGCGATTAATTTTATTTGTACAATTGTAAACATGAGAACCGAGAGTTTACCATTCCACAAATTACCTTTATTTGTATGGTCAGTGTTTTTAACAGCAATTCTTCTATTATTATCTTTACCTGTACTAGCAGGTGCTATTACAATGTTACTGACTGATAGAAATTTTAATACAACTTTTTTTGATCCAGCAGGTGGTGGTGATCCAGTTCTTTACCAGCATTTGTTT |
| Nitzschia_682 | TCTTTCTGGTATTATTGCGCACTCGGGAGGTTCTGTTGATTTAGCAATTTTCAGTCTTCATTTATCAGGAGCAGCATCTATTTTAGGAGCAATTAACTTTATTTGTACTATTGTGAATATGCGAACTGAAAGTCTGCCGTTTCATAAGTTACCTTTATTTGTTTGGGCAATTTTTATTACTGCAATTTTATTATTATTATCACTACCAGTATTAGCAGGTGCAATTACTATGTTATTAACTGATAGAAATTTTAATACTACTTTCTTTGATCCTGCTGGTGGAGGTGATCCTGTATTATATCAGCATTTATTC |
| Nitzschia_976 | TTTATCTGGTATTATCGCACACTCTGGAGGTTCTGTAGATTTAGCAATTTTCAGTCTTCACTTATCGGGAGCAGCATCAATTCTAGGCGCAATTAATTTCATTTGTACTATTATCAATATGCGAACAGAAAGTTTACCATTCCACAAATTACCTTTATTTGTTTGGGCGGTATTCATTACTGCTATTCTACTGTTATTATCACTTCCTGTACTAGCAGGAGCAATTACTATGCTGCTAACAGATAGAAATTTTAACACTACTTTCTTTGACCCTGCAGGTGGAGGTGATCCAGTGTTATATCAGCACTTATTC |
| Nitzschia_978 | TTTATCAGGTATTATTGCACACTCTGGTGGTGCCGTTGATTTAGCTATTTTCAGTTTACACCTTTCAGGTGCTGCTTCTATTTTAGGTGCGATTAATTTTATTTGTACAATTGTAAACATGAGAACCGAGAGTTTACCATTCCACAAATTACCTTTATTTGTATGGTCAGTGTTTTTAACAGCAATTCTTCTATTATTATCTTTGCCTGTACTAGCAGGTGCTATTACAATGTTACTGACTGATAGAAATTTTAATACAACTTTTTTTGATCCAGCAGGTGGTGGTGATCCAGTTCTTTACCAGCATTTGTTT |
| Nitzschia_990 | CTTTCAGGTATAATAGCTCATTCAGGTGGTGCTGTTGATTTGGCTATTTTCAGTTTACACCTTTCAGGTGCTGCATCTATTCTAGGTGCAATTAATTTCATCTGTACTATTGTAAACATGAGAACTGAAAGTTTACCATTTCATAAGTTACCATTATTTGTATGGTCAGTTTTTTTAACAGCAATTTTATTATTACTGTCTTTACCAGTATTAGCAGGTGCTATTACTATGTTATTGACTGATAGAAATTTCAACACTACTTTTTTCGACCCAGCAGGTGGTGGTGATCCAGTTCTTTACCAGCATTTGTTT |
| Nitzschia_ba0 | TTTATCAGGTATTATTGCACATTCTGGTGGTGCTGTTGATTTAGCTATTTTTAGTTTACACCTTTCAGGTGCTGCTTCTATTTTAGGTGCGATTAATTTTATTTGTACAATTGTAAACATGAGAACTGAGAGTTTACCATTCCACAAATTACCTTTATTTGTATGGTCAGTGTTTTTAACAGCAATTCTTCTATTATTATCTTTACCTGTACTAGCAGGTGCTATTACAATGTTACTGACTGATAGAAATTTTAATACAACTTTTTTTGATCCAGCAGGTGGTGGTGATCCAGTTCTTTACCAGCATTTGTTT |
| Nitzschia_bbc | TTTATCAGGTATTATTGCACACTCTGGTGGTGCTGTTGATTTAGCTATTTTCAGTTTACACCTTTCAGGTGCTGCTTCTATTTTAGGTGCGATTAATTTTATTTGTACAATTGTAAACATGAGAACTGAGAGTTTACCATTCCACAAATTACCTTTATTTGTGTGGTCGGTGTTTTTAACAGCAATCCTTCTATTATTATCTTTACCTGTACTAGCAGGTGCTATTACAATGTTACTGACTGATAGAAATTTTAATACAACTTTTTTTGATCCAGCAGGTGGGGGTGATCCAGTTCTTTACCAGCATTTGTTT |
| Nitzschia_bc0 | TTTATCAGGTATTATTGCACACTCTGGTGGTGCTGTTGATTTAGCTATTTTCAGTTTACATCTTTCAGGTGCTGCTTCTATTTTAGGTGCGATTAATTTTATTTGTACAATTGTAAACATGAGAACTGAGAGTTTACCATTCCACAAATTACCTTTATTCGTATGGTCAGTGTTTTTAACAGCAATTCTTCTATTATTATCTTTACCTGTACTAGCAGGTGCCATTACAATGTTATTGACTGATAGAAATTTTAATACAACTTTTTTTGATCCAGCAGGTGGTGGTGATCCAGTTCTTTACCAGCATTTGTTT |
| Nitzschia_c26 | TCTTTCAAGTATAACAGCGCATTCAGGAGGATCTGTGGATTTAGCGATTTTTAGTCTTCACCTATCTGGGGCTTCTTCTATTTTAGGAGCTATTAACTTTATCTGCACTATTTTTAACATGCGTACAGATAGCTTACCTTTCCACAAATTACCTTTATTTGTTTGGGCTGTTCTTATTACCGCAGTATTATTACTTTTATCTTTACCAGTTTTAGCAGGAGCTATCACAATGCTTTTAACTGATAGAAATTTTAATACAACTTTTTTTGACCCAGCTGGTGGAGGTGATCCAGTTCTTTATCAACATTTATTT |
| Nitzschia_c3f | TTTATCAGGTATTATTGCACACTCTGGTGGTGCTGTTGATTTAGCTATTTTTAGTTTACACCTTTCAGGTGCTGCTTCTATTTTAGGTGCGATTAATTTTATTTGTACAATTGTAAACATGAGAACTGAGAGTTTACCATTCCACAAATTACCTTTATTTGTATGGTCAGTGTTTTTAACAGCAATTCTTCTATTATTATCTTTACCTGTACTAGCAGGTGCTATTACAATGTTACTGACTGATAGAAATTTTAATACAACTTTTTTTGATCCAGCAGGTGGTGGTGATCCAGTTCTTTACCAGCATTTGTTT |
| Nitzschia_ceb | TTTATCAGGTATTATTGCACACTCTGGTGGTGCTGTTGATTTAGCTATTTTCAGTTTACACCTTTCAGGTGCTGCTTCTATTTTAGGTGCGATTAATTTTATTTGTACTATTGTTAACATGCGAACTGAGAGTTTACCATTCCATAAGTTACCTCTATTCGTATGGGCAGTATTTATTACCGCAATTTTATTGTTATTATCGTTACCAGTATTAGCAGGTGCAATTACCATGTTATTAACTGATAGAAATTTCAATACTACTTTCTTTGATCCAGCAGGTGGAGGTGATCCTGTATTGTATCAGCACTTATTC |
| Nitzschia_dca | TTTATCAGGTATTATTGCACACTCTGGTGGTGCTGTTGATTTAGCTATTTTCAGTTTACACCTTTCAGGTGCTGCTTCTATTTTAGGTGCGATTAATTTTATTTGTACAATTGTAAACATGAGAACTGAGAGTTTACCATTTCACAAATTACCTTTATTTGTATGGTCAGTGTTTTTAACAGCAATTCTTCTATTATTATCTTTACCTGTACTAGCAGGTGCTATTACAATGTTACTGACTGATAGAAATTTTAATACAACTTTTTTTGATCCAGCAGGTGGTGGTGATCCAGTTCTTTACCAGCATTTGTTT |
| Nitzschia_de4 | TTTATCTGGTATTATTGCACACTCTGGAGGTTCTGTAGATTTAGCAATCTTCAGTCTTCACTTATCAGGAGCGGCATCAATTCTAGGAGCAATTAATTTTATTTGTACTATTATTAATATGCGAACAGAAAGTCTGCCATTCCATAAGTTACCTTTATTCGTTTGGGCAGTATTTATTACCGCAATTTTATTATTATTATCACTACCAGTTCTAGCGGGAGCAATTACTATGTTATTAACTGATAGAAATTTCAACACCACCTTCTTTGACCCTGCGGGTGGAGGTGATCCAGTGTTATATCAGCATTTATTC |
| Nitzschia_e7b | GTTATCAGGTGTTATAGCTCACTCTGGAGGTTCTGTAGATTTAGCAATTTTCAGCCTTCATTTATCTGGAGCTGCATCTATTTTAGGTGCAATTAATTTCATTTGTACTATTGTAAATATGCGAACAGAAAGTTTACCTTTCCATAAGTTACCTCTGTTTGTATGGTCTGTTTTCATTACTGCTATTTTACTCTTATTATCTTTACCCGTATTGGCAGGAGCGATTACAATGTTACTTACAGATAGAAATTTCAATACCACTTTTTTTGATCCAGCAGGTGGTGGCGATCCGGTATTATTCCAACATTTATTC |
| Nitzschia_e95 | TCTTTCAAGTATAACAGCGCATTCAGGAGGATCTGTGGATTTAGCTATTTTTAGTCTTCACCTATCTGGGGCTTCTTCTATTTTAGGAGCTATTAACTTTATTTGCACTATTTTTAACATGCGTACAGATAGCTTACCTTTCCACAAATTACCTTTATTTGTTTGGGCTGTTCTTATTACCGCAGTATTATTACTTTTATCTTTACCAGTTTTAGCAGGAGCTATCACAATGCTTTTAACTGATAGAAATTTTAATACAACTTTTTTTGACCCAGCTGGTGGAGGTGATCCAGTTCTTTATCAACATTTATTT |
| Nitzschia_ead | TTTATCAGGCATTATTGCACACTCTGGTGGTGCTGTTGATTTAGCTATTTTCAGTTTACACCTTTCAGGTGCTGCTTCTATTTTAGGTGCGATTAATTTTATTTGTACAATTGTAAACATGAGAACTGAGAGTTTACCATTCCACAAATTACCTTTATTTGTATGGTCAGTGTTTTTAACAGCAATTCTTCTATTATTATCTTTACCTGTACTAGCAGGTGCTATTACAATGTTACTGACTGATAGAAATTTTAATACAACTTTTTTTGATCCAGCAGGTGGTGGTGATCCAGTTCTTTACCAGCATTTGTTT |
| Nitzschia_ee5 | TTTATCAGGTATTATTGCACACTCTGGTGGTGCTGTTGATTTAGCTATTTTCAGTTTACATCTTTCAGGTGCTGCTTCTATTTTAGGTGCGATTAATTTTATTTGTACAATTGTAAACATGAGAACTGAGAGTTTACCATTCCACAAATTACCTTTATTTGTATGGTCAGTGTTTTTAACAGCAATTCTTCTATTATTATCTTTACCTGTACTAGCAGGTGCCATTACAATGTTATTGACTGATAGAAATTTTAATACAACTTTTTTTGATCCAGCAGGTGGTGGTGATCCAGTTCTTTACCAGCATTTGTTT |
| Nitzschia_fce | TCTTTCAGGTATTATTGCTCACTCTGGTGGTGCAGTTGATTTAGCTATTTTTAGCTTACACCTTTCAGGCGCTGCTTCTATTTTAGGTGCAATTAATTTCATTTGTACTATTGTAAATATGAGAACTGAAAGTTTACCATTCCACAAACTTCCTTTATTTGTATGGTCAGTATTTTTGACAGCTATTTTACTGTTATTGTCTTTACCGGTACTAGCGGGCGCTATTACAATGCTGTTAACAGATAGAAATTTCAATACAACCTTTTTTGATCCTGCAGGTGGTGGAGACCCTGTTCTTTACCAACATTTATTT |
| Phaeocystis_321 | TTTGTCAGGAATTTTAGCTCATTCTGGCGGCGCTGTCGATTTAGCTATTTTTAGCTTACATTTGGCGGGTATTTCGTCTATTTTAGGAGCTATAAATTTTATTGTAACTATTTTAAACATGCGTTGTCCTGGTATGACTGCTCATAGAACACCTTTATTTGTTTGGGCTGTTTTTATAACAGCTTTTTTACTTTTGTTATCACTTCCTGTTTTAGCTGGCGCTATTACAATGTTATTAACAGATAGAAATTTTAATACGTCTTTCTTTGATCCAAACGGTGGTGGTGATCCTGTCTTATATCAGCATTTGTTT |
| Phaeocystis_4dc | GTTGTCTGGAATTTTAGCTCATTCCGGCGGTGCTGTTGATTTAGCTATTTTTAGCTTACATTTAGCTGGTATTTCTTCTATTTTGGGCGCTATAAATTTTATTGTAACGATTTTAAACATGCGTTGCCCCGGAATGGCTGCTCATAGAACACCCTTATTTGTTTGGGCTGTTTTTATTACAGCTTTTCTTCTTTTGCTCTCTCTTCCTGTTTTAGCTGGTGCTATTACTATGTTGTTAACCGATAGAAATTTTAATACTTCCTTTTTTGATCCCAGCGGTGGTGGTGATCCGGTTTTGTACCAACACTTATTT |
| Phaeocystis_8cd | GTTGTCTGGAATTTTAGCTCATTCCGGCGGTGCTGTTGATTTAGCTATTTTTAGCTTACATTTAGCTGGTATTTCTTCTATTTTGGGCGCTATAAATTTTATTGTAACTATTTTAAACATGCGTTGTCCTGGTATGACTGCTCATAGAACACCTTTATTTGTTTGGGCTGTTTTTATAACAGCTTTTTTACTTTTGTTATCACTTCCTGTTTTAGCTGGCGCTATTACAATGTTATTAACAGATAGAAATTTTAATACGTCTTTCTTTGATCCAAACGGTGGTGGTGATCCTGTCTTATATCAGCATTTGTTT |
| Phaeocystis_94b | TTTGTCAGGAATTTTAGCTCATTCTGGTGGCGCTGTCGATTTAGCTATTTTTAGCTTACATTTGGCGGGTATTTCGTCTATTTTAGGAGCTATAAATTTTATTGTAACTATTTTAAACATGCGTTGTCCTGGTATGACTGCTCATAGAACACCTTTATTTGTTTGGGCTGTTTTTATAACAGCTTTTTTACTTTTGTTATCACTTCCTGTTTTAGCTGGCGCTATTACAATGTTATTAACAGATAGAAATTTTAATACGTCTTTCTTTGATCCGAACGGTGGTGGTGATCCTGTCTTATATCAGCATTTGTTT |
| Phaeocystis_b1a | TTTGTCAGGAATTTTAGCTCATTCTGGTGGCGCTGTCGATTTAGCTATTTTTAGCTTACATTTGGCGGGTATTTCGTCTATTTTAGGAGCTATAAATTTTATTGTAACTATTTTAAACATGCGTTGTCCTGGTATGACTGCTCATAGAACACCTTTATTTGTTTGGGCTGTTTTTATAACAGCTTTTTTACTTTTGTTATCACTTCCTGTTTTAGCTGGCGCTATTACAATGTTATTAACAGATAGAAATTTTAATACGTCTTTCTTTGATCCAAACGGTGGTGGTGATCCTGTCTTATATCAGCATTTGTTT |
| Phaeocystis_c63 | TTTGTCAGGAATTTTAGCTCATTCTGGTGGCGCTGTCGATTTAGCTATTTTTAGCTTACATTTGGCGGGTATTTCGTCTATTTTAGGAGCTATAAATTTTATTGTAACTATTTTAAACATGCGTTGTCCTGGTATGACTGCTCATAGAACACCTTTATTTGTTTGGGCTGTTTTTATAACAGCTTTTTTACTTTTGTTATCACTTCCTGTTTTAGCTGGAGCTATTACAATGTTATTAACAGATAGAAATTTTAATACGTCTTTCTTTGATCCAAACGGTGGTGGTGATCCTGTCTTATATCAGCATTTGTTT |
| Prorocentrum_747 | ATTAAGCACTTCTTTCATGAGTTTATCACCTTCAAGTACAGGAAATCTTATCTTTGGATTATTAATCTCAGGTATATCCTCATGTCTCACATCTCTTAACTTTTGGACAACAATTCTAAATCTGAGCTCTTATTATCTGACATTAAAGACTATGCCATTATTCCCTTGAGCTCTCTTGATTACAGCAGGAATGCTTTTATTAACATTACCAATCTTATCAGGAGGTCTTCTAATGGTCTTGTCTGATCTTCAATCTAATACACTTTTCTTTGATCCAATCTTTGGAGGAGATCCTATATTCTATCAACACTTATTT |
| Pseudochattonella_af5 | GTTAAGTAGTGTACAGGCACATTCAGGTCCTGCCGTTGATTTAGCGATTTTTAGTTTACACTTGTCTGGAGCTTCTTCAATTTTAGGAGCTATCAACTTTATTACAACAATCCTAAATATGCGTGCACCTGGAATGAGTATGCACCGTTTACCTTTAATGGTTTGGTCGGTTTTCATTACCGCTATACTACTACTTTTATCTTTACCTGTTTTAGCTGGTGCAATTACCATGCTTTTAACAGATAGAAACTTTAACACAACATTTTTTGATCCTGCTGGAGGAGGAGATCCTGTATTATATCAGCACCTTTTC |
| Pseudochattonella_f24 | GTTAAGTAGTGTACAGGCGCATTCAGGTCCTGCCGTTGATTTAGCGATTTTTAGTTTACACTTGTCTGGAGCTTCTTCAATTTTAGGAGCTATCAACTTTATTACAACAATCCTAAATATGCGTGCACCTGGAATGAGTATGCACCGTTTACCTTTAATGGTCTGGTCGGTTTTCATTACCGCTATATTACTACTTTTATCTTTACCTGTTTTAGCTGGTGCAATTACCATGCTTTTAACAGATAGAAACTTTAACACAACATTCTTTGATCCGGCTGGAGGAGGAGATCCTGTACTATATCAGCACCTTTTC |
| Pseudonitzschia_220 | TCTTTCAGGAGTTTTAGCTCATTCAGGAGGTTCTGTTGATTTGGCAATTTTCAGTCTTCACTTATCAGGTGCTGCGTCAATTTTAGGTGCAATTAATTTTATTTGTACTATTGTTAATATGCGAACTGAAAGTTTACCATTCCATAAACTTCCTTTATTTGTTTGGTCAGTGTTTATTACTGCGATCTTGTTACTGTTATCACTACCAGTTTTGGCAGGAGCAATTACAATGTTGTTAACAGATAGAAATTTTAACACAACATTTTTTGATCCAGCAGGTGGTGGAGATCCTGTCCTTTTTCAGCATTTATTT |
| Pseudonitzschia_4e5 | ACTTTCAGGAGTTTTATCTCACTCAGGAGGGTCTGTTGATTTAGCAATTTTCAGTCTTCATTTATCAGGAGCAGCATCAATTTTAGGTGCAATTAATTTTATTTGTACAATTGTAAATATGCGAACAGAGAGTTTACCATTTCATAAACTTCCTTTATTCGTTTGGGCTGTTTTTATTACAGCTATTTTACTGTTACTATCGTTACCTGTTTTAGCAGGAGCGATTACAATGTTATTAACAGATAGAAATTTTAATACTACTTTCTTTGATCCAGCAGGTGGAGGAGATCCAGTTCTTTTCCAGCATTTGTTT |
| Pseudonitzschia_95f | TCTTTCTGGAGCTATTGCTCATTCTGGAGGGTCTGTCGATCTAGCAATTTTTAGTCTTCATTTATCTGGAGCTGCATCAATTTTAGGTGCAATTAATTTCATTTGTACCATTGTAAATATGCGAACTGAAAGTTTACCTTTCCACAAACTTCCTTTATTTGTGTGGGCTGTTTTCATTACCGCTATTTTACTTTTATTATCTCTACCTGTACTAGCTGGAGCAATTACAATGTTGTTGACTGATCGGAACTTTAACACAACTTTTTTTGATCCAGCAGGCGGAGGAGATCCTGTGCTATTCCAGCACTTATTC |
| Pseudonitzschia_d36 | TCTTTCTGGAGTTTTATCCCACTCTGGAGGTTCTGTTGATTTAGCAATTTTCAGTCTTCATTTGTCTGGGGCAGCTTCTATTTTAGGAGCGATTAACTTCATTTGTACAATTGTAAATATGAGAACTGAAAGTTTACCATTCCACAAACTTCCTCTGTTTGTATGGGCAGTTTTCATTACTGCTATTTTATTACTTTTATCATTACCAGTTTTAGCTGGAGCAATTACAATGTTGTTAACTGACAGGAACTTTAATACTACATTTTTTGACCCTGCAGGTGGAGGTGATCCAGTACTTTTCCAACATTTATTC |
| Pseudonitzschia_d40 | TCTTTCAAGTGTTTTATCTCACTCTGGAGGTTCTGTGGATTTAGCAATTTTCAGTCTTCACTTGTCAGGAGCAGCTTCAATTTTAGGTGCAATTAATTTTATTTGTACAATTGTAAATATGAGAACTGAGAGTTTACCATTTCATAAGCTTCCGTTATTTGTTTGGTCTGTTTTCATTACTGCCATTTTATTACTACTATCTTTACCTGTTCTAGCGGGTGCTATAACAATGTTATTGACCGACAGAAATTTCAATACCACTTTTTTTGATCCTGCAGGTGGAGGAGATCCTGTACTTTTTCAGCACTTATTT |
| Pseudonitzschia_e48 | TCTTTCAGGAGTTTTATCTCATTCAGGAGGTTCTGTTGATCTAGCAATTTTTAGCCTTCATTTATCTGGGGCAGCGTCAATTTTAGGTGCGATCAATTTTATTTGTACTATTGTAAATATGAGAACTGAAAGTTTACCATTCCACAAACTTCCTTTATTTGTTTGGTCTGTTTTTATTACAGCCATTTTATTATTATTAGCACTACCTGTTTTAGCAGGAGCAATTACAATGTTATTAACTGATAGAAATTTCAATACTACCTTTTTTGATCCCGCAGGTGGAGGAGATCCTGTACTTTTTCAACATTTGTTT |
| Woloszynskia_584 | ATTATCTACTTCTTTTATGACTTTATCACCTTCAAGTACAGGAAATCTTATCTTTGGATTATTAATCTCTGGTATATCCTCATGTCTTACATCTCTTAACTTTTGGATTACAATTCTAAATCTGAGATCTTATTATCTGACATTAAAGACTATCCCATTATTTCCTTGGGCTTTCTTGATTACAGCTTTCATGCTTTTATTAACATTACCAATTTTATCTGGTACACTTATTTTAATATTAGGTGATCTTCATTCTAATACACTTTTCTTTGATCCAATATTTGGAGGAGATCCTATATTCTATCAACACTTATTT |

Table S3: Complete list of logistic-regression models tested describing taxon occurrence as a function of sea-surface temperature, pH, and salinity.

| Taxon | Models |
| --- | --- |
| Alexandrium_2b2 | (1 + pHStd \| Season), (1 + TempStd \| Season), (1 + SalinityStd \| Season), TempStd + (1 + pHStd \| Season), pHStd + (1 + TempStd \| Season), pHStd + SalinityStd + (1 + TempStd \| Season), pHStd + TempStd, pHStd * TempStd, pHStd + SalinityStd, pHStd + SalinityStd + (1 \| Season), pHStd + (1 + SalinityStd \| Season), SalinityStd + (1 + pHStd \| Season), pHStd + (0 + SalinityStd \| Season), SalinityStd + (0 + pHStd \| Season), 0 + pHStd + (1 + TempStd \| Season), pHStd + (0 + TempStd \| Season) |
| Alexandrium_3fc | (1 + pHStd \| Season), (1 + TempStd \| Season), (1 + SalinityStd \| Season), TempStd + (1 + pHStd \| Season), pHStd + (1 + TempStd \| Season), pHStd + SalinityStd + (1 + TempStd \| Season), pHStd + TempStd, pHStd * TempStd, pHStd + SalinityStd, pHStd + SalinityStd + (1 \| Season), pHStd + (1 + SalinityStd \| Season), SalinityStd + (1 + pHStd \| Season), pHStd + (0 + SalinityStd \| Season), SalinityStd + (0 + pHStd \| Season), 0 + (1 + TempStd \| Season), 1 + (0 + TempStd \| Season) |
| Hematodinium_449 | (1 + pHStd \| Season), (1 + TempStd \| Season), (1 + SalinityStd \| Season), TempStd + (1 + pHStd \| Season), pHStd + (1 + TempStd \| Season), pHStd + SalinityStd + (1 + TempStd \| Season), pHStd + TempStd, pHStd * TempStd, pHStd + SalinityStd, pHStd + SalinityStd + (1 \| Season), pHStd + (1 + SalinityStd \| Season), SalinityStd + (1 + pHStd \| Season), pHStd + (0 + SalinityStd \| Season), SalinityStd + (0 + pHStd \| Season) |
| Karlodinium_8ed | (1 + pHStd \| Season), (1 + TempStd \| Season), (1 + SalinityStd \| Season), TempStd + (1 + pHStd \| Season), pHStd + (1 + TempStd \| Season), pHStd + SalinityStd + (1 + TempStd \| Season), pHStd + TempStd, pHStd * TempStd, pHStd + SalinityStd, pHStd + SalinityStd + (1 \| Season), pHStd + (1 + SalinityStd \| Season), SalinityStd + (1 + pHStd \| Season), pHStd + (0 + SalinityStd \| Season), SalinityStd + (0 + pHStd \| Season), 0 + (1 + TempStd \| Season), 1 + (0 + TempStd \| Season), 0 + (0 + TempStd \| Season) |
| Karlodinium_a27 | (1 + pHStd \| Season), (1 + TempStd \| Season), (1 + SalinityStd \| Season), TempStd + (1 + pHStd \| Season), pHStd + (1 + TempStd \| Season), pHStd + SalinityStd + (1 + TempStd \| Season), pHStd + TempStd, pHStd * TempStd, pHStd + SalinityStd, pHStd + SalinityStd + (1 \| Season), pHStd + (1 + SalinityStd \| Season), SalinityStd + (1 + pHStd \| Season), pHStd + (0 + SalinityStd \| Season), SalinityStd + (0 + pHStd \| Season), 0 + pHStd + (1 + SalinityStd \| Season) |
| Pseudonitzschia_4e5 | 0 + TempStd + SalinityStd + (1 \| Season), TempStd + SalinityStd + (1 \| Season), TempStd + SalinityStd, TempStd + SalinityStd + pHStd, pHStd + TempStd + SalinityStd + (1 \| Season) |
| Pseudonitzschia_d36 | pHStd + TempStd + SalinityStd + (1 \| Season), pHStd + SalinityStd + (1 \| Season), 0 + pHStd + SalinityStd + (1 \| Season), 0 + SalinityStd + (1 \| Season), TempStd + SalinityStd + pHStd, 0 + (1 + SalinityStd \| Season), 0 + TempStd + (1 + SalinityStd \| Season), 0 + TempStd + pHStd + (1 + SalinityStd \| Season) |
| Pseudonitzschia_d40 | SalinityStd + TempStd + (1 \| Season), 1 + SalinityStd + (0 + TempStd\| Season), 1 + SalinityStd + (1 + TempStd\| Season), 0 + SalinityStd + (1 \| Season), SalinityStd |

Table S4: Top ten taxa associated with Alexandrium_2b2 by CAP, with association strength and direction.

| Taxon | CAP1 |
| --- | --- |
| Ditylum_ba4 | 0.2017 |
| Ditylum_a31 | 0.2001 |
| Thalassiosira_47a | 0.1962 |
| Poteriospumella_86a | 0.1837 |
| Oncorhynchus_584 | 0.1837 |
| Poteriospumella_b57 | -0.1634 |
| Chattonella_135 | 0.1563 |
| Balanus_513 | 0.1501 |
| Pinnularia_950 | 0.147 |
| Cylindrotheca_072 | 0.1462 |

Table S5: Top ten taxa associated with Alexandrium_3fc by CAP, with association strength and direction.

| Taxon | CAP1 |
| --- | --- |
| Prasinoderma_6ac | 0.2237 |
| Stephanodiscus_0c0 | -0.2237 |
| Chaetoceros_dcd | 0.2081 |
| Phaeocystis_4dc | -0.2081 |
| Rhizosolenia_006 | -0.1977 |
| Rhizosolenia_d78 | 0.1821 |
| Chaetoceros_734 | 0.1821 |
| Mytilus_4a8 | 0.1768 |
| Hincksia_8a1 | -0.1768 |
| Minutocellus_5ae | -0.1612 |

Table S6: Top ten taxa associated with Hematodinium_449 by CAP, with association strength and direction.

| Taxon | CAP1 |
| --- | --- |
| Saxidomus_33e | 0.2105 |
| Chrysochromulina_7aa | 0.1741 |
| Cylindrotheca_799 | 0.1579 |
| Thalassionema_f2c | -0.1457 |
| Gelidiophycus_401 | 0.1396 |
| Nitzschia_dca | -0.1376 |
| Obelia_f83 | -0.1356 |
| Karlodinium_a27 | 0.1235 |
| Balanus_2eb | 0.1174 |
| Stephanodiscus_0c0 | -0.1174 |

Table S7: Top ten taxa associated with Karlodinium_8ed by CAP, with association strength and direction.

| Taxon | CAP1 |
| --- | --- |
| Ditylum_a31 | 0.2405 |
| Thalassiosira_6b3 | 0.2164 |
| Poteriospumella_b57 | -0.1924 |
| Ditylum_ba4 | 0.1804 |
| Cylindrotheca_942 | -0.1683 |
| Cylindrotheca_072 | 0.1683 |
| Oncorhynchus_584 | 0.1683 |
| Pythium_482 | 0.1683 |
| Pythium_67b | 0.1683 |
| Asterionellopsis_681 | 0.1563 |

Table S8: Top ten taxa associated with Karlodinium_a27 by CAP, with association strength and direction.

| Taxon | CAP1 |
| --- | --- |
| Balanus_2eb | -0.1727 |
| Hincksia_8a1 | -0.1507 |
| Skeletonema_0a6 | -0.1369 |
| Gelidiophycus_401 | -0.1357 |
| Micromonas_6f9 | -0.1313 |
| Pythium_3b3 | -0.12 |
| Hematodinium_449 | -0.1149 |
| Balanus_6ef | -0.1131 |
| Bathycoccus_14e | -0.1093 |
| Pycnococcus_d91 | -0.1093 |

Table S9: Top ten taxa associated with Pseudonitzschia_4e5 by CAP, with association strength and direction.

| Taxon | CAP1 |
| --- | --- |
| Calanus_79b | 0.2233 |
| Pseudonitzschia_d36 | 0.1932 |
| Rhizosolenia_e51 | 0.1932 |
| Ditylum_ce4 | 0.1751 |
| Clytia_8b4 | 0.1751 |
| Peronospora_8ec | 0.169 |
| Chaetoceros_734 | 0.169 |
| Pseudocalanus_175 | 0.163 |
| Asterionellopsis_fce | 0.1569 |
| Dendraster_0f8 | 0.1509 |

Table S10: Top ten taxa associated with Pseudonitzschia_d36 by CAP, with association strength and direction.

| Taxon | CAP1 |
| --- | --- |
| Calanus_79b | 0.229 |
| Clytia_8b4 | 0.1925 |
| Pseudonitzschia_4e5 | 0.1783 |
| Asterionellopsis_fce | 0.1739 |
| Asterionellopsis_681 | 0.1677 |
| Dendraster_0f8 | 0.1677 |
| Clytia_cae | 0.1535 |
| Skeletonema_cf5 | 0.1516 |
| Pseudocalanus_175 | 0.1411 |
| Chondracanthus_0d6 | 0.1331 |

Table S11: Top ten taxa associated with Pseudonitzschia_d40 by CAP, with association strength and direction.

| Taxon | CAP1 |
| --- | --- |
| Ditylum_ce4 | 0.2462 |
| Synchaeta_06c | 0.1977 |
| Calanus_79b | 0.1916 |
| Skeletonema_0a6 | 0.1916 |
| Pseudocalanus_175 | 0.1882 |
| Attheya_223 | 0.1766 |
| Chaetoceros_00e | 0.1712 |
| Thalassiosira_391 | 0.1616 |
| Ditylum_a31 | 0.1541 |
| Pseudonitzschia_4e5 | 0.1473 |
